# Decoding the RNA Splicing Network in HNRNPH2-R114W Brain Organoids

**DOI:** 10.64898/2026.02.06.704442

**Authors:** Carlo Donato Caiaffa, Neda Ghousifam, Stephan Lloyd Watkins, Helder I. Nakaya, Rodney Bowling, Richard H. Finnell, Robert M. Cabrera

## Abstract

Autism spectrum disorder and related neurodevelopmental diseases are increasingly linked to disrupted RNA processing during corticogenesis, yet resolving variant-specific mechanisms during early human development remains challenging. We generated isogenic cerebral organoids from a patient iPSC line carrying the HNRNPH2-R114W mutation and its CRISPR-corrected counterpart. Developmental validation by time-course imaging, followed by bulk RNA-Seq at day 20, showed comparable early differentiation and corticogenesis. Genotype was the primary determinant of global expression state, with organized remodeling of neuronal and metabolic programs. Splicing analyses demonstrated junction-dominant dysregulation, where 6,310 junctions exhibited significant differential usage. Junction-level resolution revealed a dominant genotype-related signal than exon-centric analysis. A structural docking model shows HNRNPH2 RNA recognition motifs 1-3 within a spliceosomal context and position residue R114 at an RNA-contact cleft, consistent with direct disruption of RNA engagement by R114W. Integrating differential expression with event- and isoform-level evidence prioritized convergent regulatory hubs enriched for axon guidance, synaptic signaling, and extracellular matrix programs. Single-cell RNA-Seq deconvolution revealed modest compositional shifts that do not account for the magnitude of transcriptional and splicing alterations. Collectively, these data support that HNRNPH2-R114W drives pervasive, junction-centered RNA rewiring in human cortical organoids and defines isoform-level endpoints for mechanistic studies and therapeutic testing in an isogenic developmental model.

## INTRODUCTION

Autism spectrum disorder (ASD) and related neurodevelopmental diseases (NDDs) arise from diverse genetic and environmental factors that converge on early cortical development, including neuronal differentiation, circuit assembly, and synaptic maturation. Human genetic and transcriptomic studies increasingly implicate RNA processing, particularly alternative splicing, as a key mediator of ASD/NDD risk, reflecting the highly dynamic splicing programs that accompany corticogenesis and neuronal subtype specification (Raj & Blencowe, 2015; Li et al., 2016; Weyn-Vanhentenryck et al., 2018). The pathogenic trajectory from mutation to ASD reflects isoform-level alterations in critical neurodevelopmental genes rather than changes in total gene expression, motivating the development of experimental systems and analytical frameworks that explicitly resolve transcript and splice-junction structures.

Splicing regulation in the developing brain functions as a dynamically coordinated and programmable network, rather than a static housekeeping process. This intricate regulation ensures the production of neuron-specific isoform repertoires, modulates neuronal excitability, and shapes synaptic signaling during defined developmental windows (Raj & Blencowe, 2015; Ule & Blencowe, 2019). Throughout fetal corticogenesis, RNA-binding proteins (RBPs) guide temporally precise splicing transitions. Disruption of these RBP networks results in widespread yet interpretable changes, which preferentially impact synaptic and projection-related gene sets that are consistently enriched among individuals with ASD and NDDs (Huelga et al., 2012; Irimia et al., 2014; Li et al., 2016; Weyn-Vanhentenryck et al., 2018).

ASD-associated misregulation of conserved neuronal microexons, which are enriched in synaptic genes and tightly regulated during cortical development, exemplifies how subtle splicing defects can destabilize synaptic maturation and circuit function (Irimia et al., 2014; Raj & Blencowe, 2015). Integrative genetic and transcriptomic analyses further show that splicing quantitative trait loci and isoform shifts explain a substantial fraction of complex trait associations, reinforcing splicing as a major conduit by which genetic variation influences neurodevelopmental phenotypes (Li et al., 2016). These observations argue that both gene-level expression and exon–junction architectures need to be interrogated to understand ASD-linked RNA dysregulation, directly motivating an approach that integrates differential expression and splice-junction analyses.

Neuron-enriched RBPs such as RBFOX and NOVA, together with heterogeneous nuclear ribonucleoproteins (HNRNPs), organize regulatory splicing maps that define neuronal isoform landscapes in the context of long introns and complex exon architectures (Raj & Blencowe, 2015; Ule & Blencowe, 2019). Within this framework, HNRNP H-family proteins act as key modulators of splice-site selection and suppressors of aberrant junctions at G-rich motifs, suggesting that their disruption will preferentially manifest as junction-centric remodeling of neuronal isoform programs rather than simple exon gain or loss (Honoré et al., 1995; Caputi & Zahler, 2001; Huelga et al., 2012; Ule & Blencowe, 2019).

Human induced pluripotent stem cell (iPSC) - derived brain organoids capture key features of early corticogenesis, including ventricular zone-like progenitors, cortical plate-like neurons, and emerging network activity, in a system that is sufficiently reproducible to support modeling of monogenic NDDs (Lancaster et al., 2013; Pașca et al., 2015; Mariani et al., 2015; Quadrato et al., 2017; Bershteyn et al., 2017; Velasco et al., 2019). For RNA-processing genes such as splicing regulators and HNRNP proteins, organoids provide the appropriate cell types and developmental timing to interpret gene-level expression changes together with junction- and isoform-level splicing outcomes.

Bulk organoid transcriptomes are influenced by maturation state and cell-type composition, which can differ subtly between lines and batches, thereby confounding regulatory signatures (Qian et al., 2016; Quadrato et al., 2017; Velasco et al., 2019). Deconvolution approaches that project bulk RNA-seq data onto multi-subject single-cell reference atlases, such as MuSiC (multi-subject single cell deconvolution approach) integrated with a single-cell Seq-based human neural organoid atlas (HNOCA) (Wang et al., 2019; He et al., 2024), enable estimation of cell-type fractions and improve interpretation of expression and splicing changes by distinguishing cell-intrinsic effects from shifts in lineage balance (Nowakowski et al., 2017; Wang et al., 2019; Pollen et al., 2019; He et al., 2024).

To strengthen causal inference in this setting, genome-edited isogenic pairs, where a patient HNRNPH2-R114W rare variant is corrected on a matched genetic background, minimize inter-individual variability and are particularly valuable for RNA-processing genes with extensive downstream effects (Qian et al., 2016; Bershteyn et al., 2017; Velasco et al., 2019; He et al., 2024). Such isogenic organoid systems increase the power to detect genotype-linked changes in expression, splice-junction usage, and inferred cell-type composition, and to assess whether correction of a single RBP variant rescues multi-layer RNA phenotypes.

Rare pathogenic variants in HNRNPH2 are of particular interest because they directly affect the RNA-processing machinery and cause an X-linked neurodevelopmental syndrome characterized by developmental delay, seizures, and frequent autistic traits (Bain et al., 2016; Pilch et al., 2018; Gillentine et al., 2021). HNRNPH2 encodes an HNRNP H-family protein with quasi-RNA recognition motifs and a PY-type nuclear localization signal that mediates binding to G-rich RNA elements and nuclear import via the karyopherin-β2 pathway (Honoré et al., 1995; Caputi & Zahler, 2001; Brownmiller & Caplen, 2023). Missense variants cluster in the PY-NLS and qRRM domains are predicted to impair nuclear trafficking and alter RNA-binding specificity, leading to widespread changes in splice-site selection and junction usage at HNRNP H/F target motifs (Caputi & Zahler, 2001; Huelga et al., 2012; Conboy, 2017; Brownmiller & Caplen, 2023). Patient-derived models show deregulation of hundreds of genes and extensive alternative splicing, supporting dominant-negative or toxic gain-of-function mechanisms with limited compensation by the paralog HNRNPH1 (Gillentine et al., 2021; Larizza et al., 2022; Brownmiller & Caplen, 2023).

Family-level analyses of HNRNP genes indicate that deleterious variants across this group produce partially overlapping NDD phenotypes, giving rise to the concept of ‘‘RBPopathy”, in which disruption of RBP networks represents a recurrent mechanism underlying ASD/ID (Gillentine et al., 2021; Brownmiller & Caplen, 2023). Within this framework, HNRNPH2 sits in splicing regulatory networks enriched for synaptic genes and ASD risk loci and is expected to perturb isoform programs linked to excitatory-inhibitory balance and synaptic maturation when disrupted (Irimia et al., 2014; Raj & Blencowe, 2015; Brownmiller & Caplen, 2023).

Organoid-based ASD models demonstrate that early neurodevelopmental processes, including progenitor dynamics, neuronal differentiation, and nascent network formation, can be altered by risk variants without overt morphological abnormalities, consistent with the clinical presentation of HNRNPH2-related NDD (Lancaster et al., 2013; Mariani et al., 2015; Quadrato et al., 2017; Bershteyn et al., 2017). Together with genetic and biochemical evidence, these observations support a model in which HNRNPH2-linked ASD/ID represents an RBP-driven spliceopathy, in which instability of splicing networks during cortical development is a central pathogenic mechanism (Irimia et al., 2014; Raj & Blencowe, 2015; Brownmiller & Caplen, 2023).

Given this background, a multi-layer RNA-centric approach is required to elucidate how pathogenic HNRNPH2 variants alter human cortical transcriptomes. Gene-level differential expression analysis provides a foundation for identifying genotype-linked transcriptional changes and define global remodeling in isogenic cerebral organoids (Robinson et al., 2010; Love et al., 2014). Because HNRNPH2 primarily regulates splicing, junction- and feature-centric methods are necessary to capture splice-junction architecture and isoform rebalancing that may be hidden at the exon or gene level (Pervouchine et al., 2013; Goldstein et al., 2016). In parallel, cell-type deconvolution of bulk organoid RNA-seq using multi-subject single-cell cortical reference atlases, such as MuSiC (Wang et al., 2019; Nowakowski et al., 2017; Pollen et al., 2019), helps distinguish within-cell-type regulatory changes from modest shifts in lineage proportions.

Despite strong clinical and mechanistic evidence implicating HNRNPH2 in NDD and ASD, the causal chain from an HNRNPH2 variant to human cortical RNA phenotypes remains incompletely resolved in tissue-relevant contexts where splicing programs and RBP networks are active (Lancaster et al., 2013; Qian et al., 2016; Velasco et al., 2019; Wang et al., 2019). Here, an isogenic pair of HNRNPH2-R114W rare variant and CRISPR-corrected cerebral organoids provides a human developmental system in which gene-level expression, isoform- and junction-level splicing, and inferred cell-type composition can be quantified within a matched genetic background. In this study, these multi-layer readouts are integrated to define how the HNRNPH2-R114W variant reprograms human cortical transcriptomes, to delineate junction-dominant spliceopathy and its relationship to differential expression, and to identify isoform-level endpoints that may inform precise therapeutic strategies for ASD-linked RBP dysregulation.

## RESULTS

### Validation of iPSC lines and maintenance of pluripotency

To objectively assess the functional readouts of the HNRNPH2-R114W mutation during early neuronal development, isogenic cerebral organoids were generated from patient-derived PBMCs using HNRNPH2-R114W rare variant (Minus) and CRISPR-corrected (Plus) iPSC lines. Both Minus and Plus iPSC lines displayed robust nuclear POU5F1 (Oct4) and NANOG expression (Fig. 1A; Supplemental Fig. 1 A-B), confirming the maintenance of pluripotency in both genotypes under standard culture conditions. Additionally, HNRNPH2 signal intensity was moderately reduced in Minus, while SHANK1 demonstrated a modest increase and substantially greater variability (Fig. 1A; Supplemental Fig. 1 E-F).

**Figure 1.**
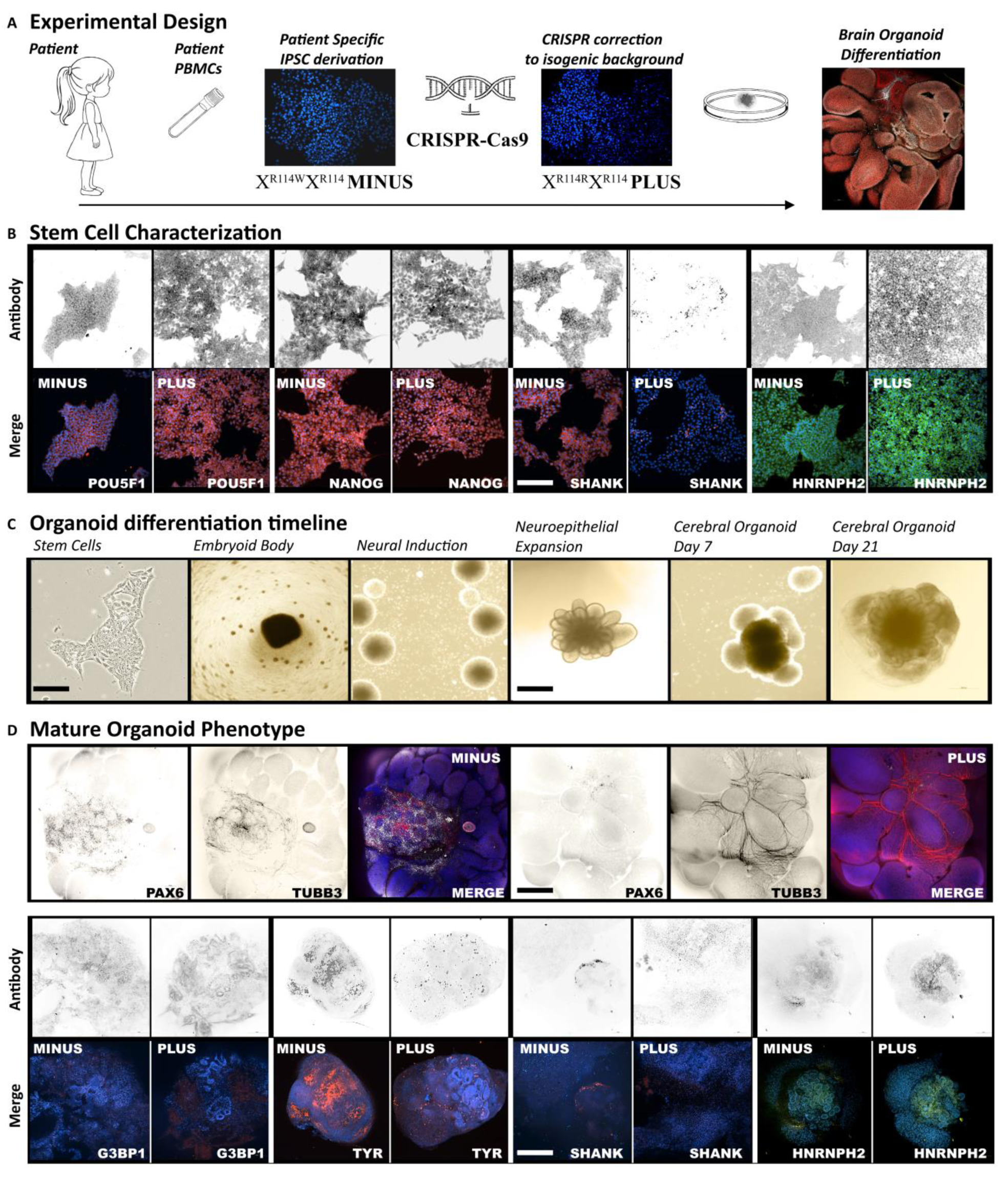
Characterization of stem cells and cerebral organoid differentiation establishes an isogenic HNRNPH2-R114W model that maintains early neurodevelopmental architecture. (A) Schematic overview of the experimental workflow, illustrating the generation of patient PBMC-derived iPSC lines harboring the HNRNPH2 R114W variant (“Minus”) and the CRISPR-corrected isogenic control (“Plus”), followed by differentiation into cerebral organoids. (B) Immunofluorescence analysis verifies pluripotency in both Minus and Plus iPSCs using nuclear POU5F1 (OCT4) and NANOG, alongside parallel staining for HNRNPH2 and SHANK. The scale bar represents 200 µm (C) Timeline schematic of the cerebral organoid protocol, detailing progression from stem cells through embryoid body formation, neural induction, and organoid maturation, with representative brightfield images at key stages, including days 7 and 21. The scale bar represents 100 and 200 µm, respectively (D) Representative immunostaining of organoids demonstrates neuroepithelial and neuronal markers (PAX6 and TUBB3) with merged views, as well as additional marker panels comparing Minus and Plus organoids. The scale bar represents 100 µm.

### Organoid differentiation timeline and morphological development

The cerebral organoid differentiation protocol followed a standardized timeline encompassing embryoid body formation, neural induction, neuroepithelial expansion, and early organoid maturation. At each corresponding stage, both Minus and Plus lineages generated neuroepithelial structures with central lumens and radially organized progenitor zones, ultimately forming larger, laminated neuroepithelial domains without discernible morphological abnormalities. Notably, organoids derived from both genotypes exhibited comparable size and gross morphology at matched time points, with no overt morphological differences observed (Fig. 1B).

### Preservation of neuroepithelial and neuronal identity in the context of the HNRNPH2-R114W mutation

Whole-mount immunostaining confirmed the expression of canonical neurodevelopmental markers. PAX6-positive cells formed radial, ventricular-like zones surrounding central luminal spaces, while TUBB3-positive neurons localized to peripheral regions in both genotypes. Merged channel analysis revealed an interconnected distribution of PAX6 and TUBB3, consistent with organoid maturation. No significant reduction of PAX6 domains or depletion of TUBB3-positive neurons was observed in Minus compared to Plus organoids, suggesting that neuroepithelial and neuronal identities are largely preserved (Fig. 1C; Supplemental Fig. 1 G-H).

### HNRNPH2 immunodetection reveals genotype-dependent distribution patterns without significant differences in overall abundance

HNRNPH2 immunostaining at the protein level demonstrated genotype-dependent distribution patterns, although overall detection remained similar (Fig. 1C; Supplemental Fig. 1 I). In Plus organoids, HNRNPH2 was predominantly localized within neuroepithelial compartments, while Minus organoids exhibited weaker and more diffuse HNRNPH2 staining throughout the tissue, suggesting altered cellular localization associated with the mutation. By day 30, G3BP1 identified stress-responsive RNA granule structures, and TYR-positive cells delineated pigmented regions enriched in the Minus lineage, consistent with melanocyte-like cell populations observed in subsequent RNA analyses. SHANK1 marked synaptic candidate regions in both genotypes, with a pronounced increase in Plus organoids (Supplemental Fig. 1 J-L). Together, these findings validate the differentiation of human iPSC-derived organoids and establish their biological suitability for transcriptomic analysis, linking morphology and marker expression to downstream analyses.

### Global transcriptional remodeling in HNRNPH2-R114W organoids

The gene expression patterns of Minus and Plus organoids were compared using a DESeq2 pipeline (Fig. 2A). Pearson correlation showed high within-group similarity and clear separation between groups (Fig. 2B). Additional ordination analyses confirmed the genotype-based separation, with PCA primarily distinguishing Minus and Plus along PC1, the dominant axis capturing the largest source of variation in the dataset, explaining 99.5% of the variance, while PC2 explained only 0.3% (Fig. 2C; Supplemental Fig. 2B).

**Figure 2.**
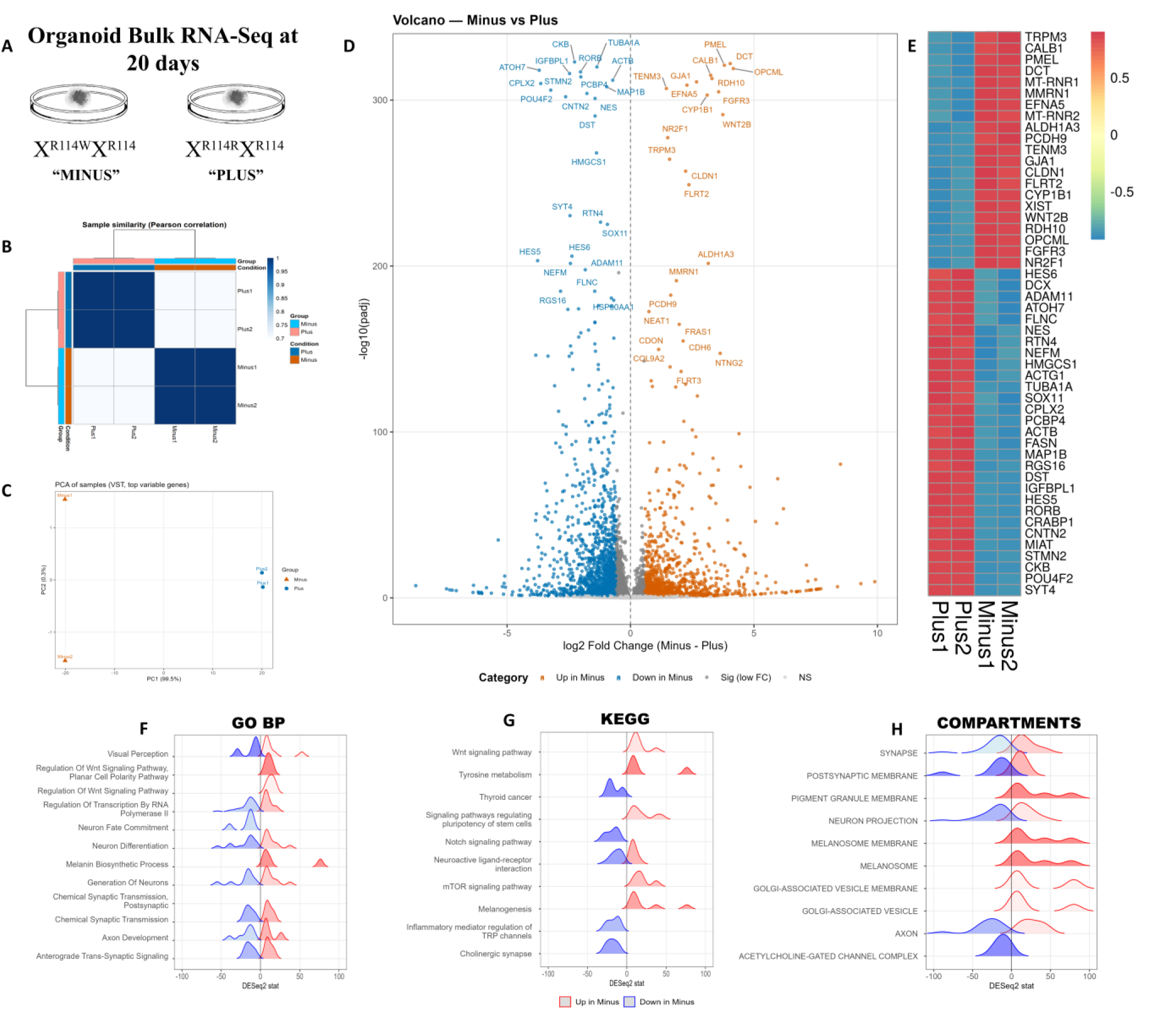
HNRNPH2-R114W organoids display a genotype-specific transcriptomic shift as determined by bulk RNA sequencing. (A) Schematic of the study design, indicating bulk RNA-seq profiling of Minus and Plus organoids at day 20. (B) Sample-to-sample similarity is evaluated using a Pearson correlation heatmap computed on a variance-stabilized (VST) expression matrix of the top 500 most variable genes, demonstrating within-genotype concordance and between-genotype separation. (C) The global sample structure is visualized using principal component analysis (PCA) based on the same VST matrix, highlighting genotype-driven separation. (D) Differential expression is depicted in a volcano plot for Minus versus Plus organoids using DESeq2, with effect size (log2 fold change) and significance (BH-adjusted p-value); significant genes are defined by padj < 0.05 and baseMean ≥ 10, with log2 fold-change shrinkage applied for visualization where indicated. (E) Heatmap displays the top differentially expressed genes across all samples on the VST scale. (F, G, H) Functional enrichment analyses for genotype-associated expression changes are presented as ridge or peak-density panels for GO Biological Process, subcellular compartment annotations, and KEGG pathways, respectively. Gene-level preprocessing includes removing genes with fewer than 10 total reads and collapsing duplicate gene identifiers after Ensembl version stripping.

Effect size and statistical significance were graphed as a volcano plot (Fig. 2D; Supplemental Fig. 2E), representing a large set of differentially expressed genes with broadly similar numbers of up and downregulated transcripts at FDR 0.05 and |log2FC| ≥ 0.58. A heatmap of the top 50 differentially expressed genes also showed clear signature patterns (Fig. 2E; Supplemental Fig. 2D), including genes that control both neurodevelopment and pigment pathways. Top genes, like CNTN2, TUBA1A, CRABP1, and RDH10, were also identified as key differential splicing targets, suggesting an overlap between gene expression and splicing (Fig. 3).

**Figure 3.**
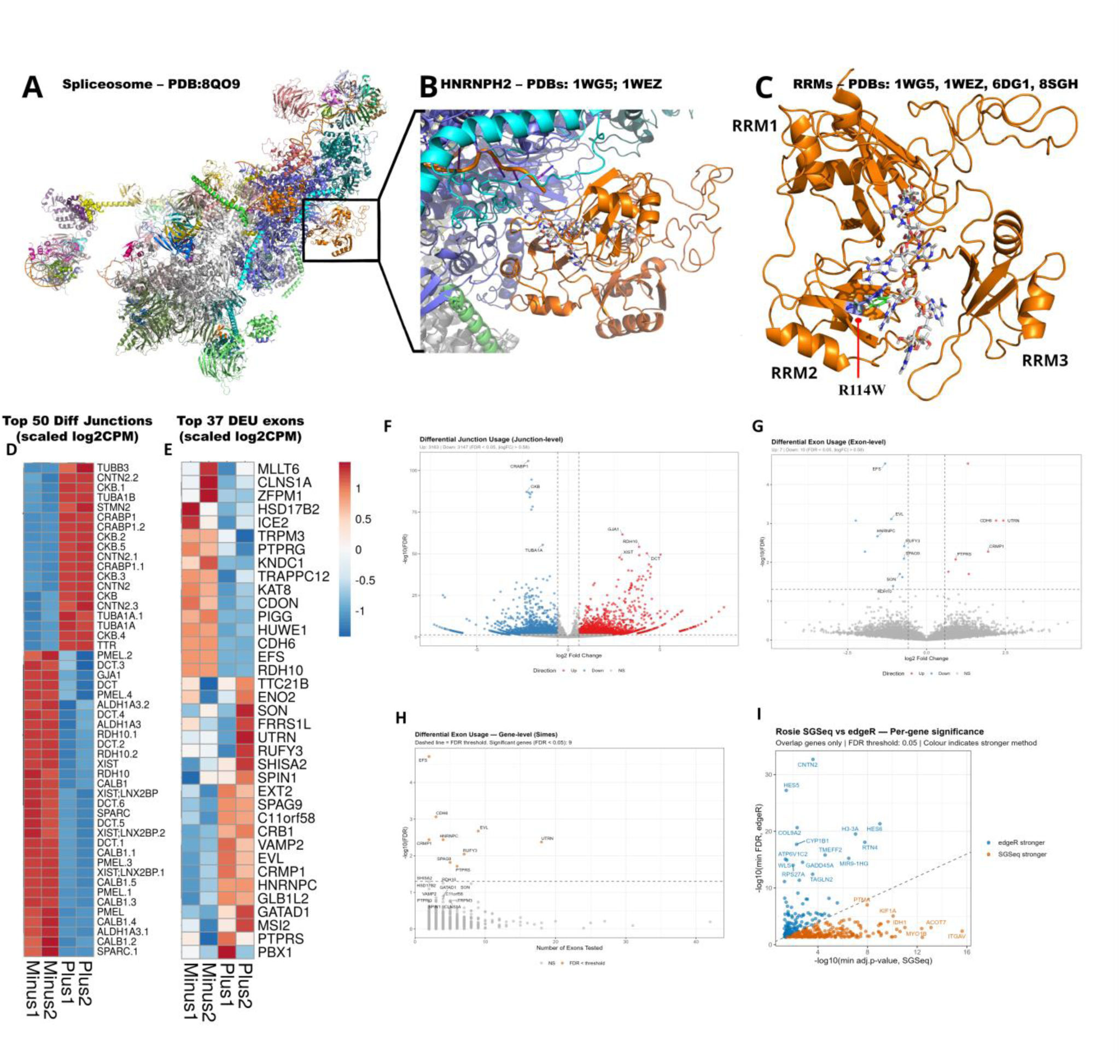
Splicing dysregulation in HNRNPH2-R114W organoids is predominantly junction-driven and only partially detected by exon-centric analyses. (A) Structural docking model depicts HNRNPH2 within the spliceosomal context (spliceosome PDB 8QO9), shown as an orange cartoon with the docking region boxed. (B) Enlarged view of the boxed region from (A) illustrates the inferred HNRNPH2–spliceosome interface and the continuity of the HNRNPH2-associated RNA path into spliceosomal RNA. (C) Structural model assembled from PDB entries 1WGS, 1WEZ, 2LEB, 4C0O, and 6HPJ highlights residue 114 at the RNA-binding cleft (R114 highlighted; W114 mutation overlaid), exhibiting a direct RNA-contact mechanism for the R114W substitution. (D) Heatmap displays the top 50 differentially used splice junctions across Minus and Plus samples (logCPM/log2CPM scale with prior count; row-scaled for visualization). (E) The heatmap shows the top differentially used exons across samples on the same scale. (F) Genome-wide junction-centric differential usage results are derived from junction discovery and quantification, including annotated and novel junctions supported by split reads, and tested with limma-trend and empirical Bayes moderation; significance is defined at BH FDR < 0.05, with direction labels optionally using |log2FC| ≥ 0.58 for plotting. (G) Exon-centric differential usage results are based on exon counts generated with featureCounts using strand-specific counting and multimappers excluded, normalized by TMM, filtered to retain exons with at least 5 counts in at least 2 samples, and tested using voom and diffSplice, where significance is defined at BH FDR < 0.05. (H) Gene-level aggregation of exon-centric splicing signals is performed using a Simes framework to summarize differential exon usage by gene. (I) The Concordance overview compares gene-level splicing signals across junction-centric and exon/feature-centric workflows, highlighting both overlaps and method-specific detections.

Functional enrichment analysis across GO, Reactome, Compartments, KEGG, and Hallmark annotations (Fig. 2F–H; Supplemental Fig. 2G-I) showed that genes downregulated in Minus were enriched for neurodevelopmental processes, including neuron differentiation, axon development, and synaptic transmission. In contrast, genes upregulated in Minus were enriched for WNT/Notch signaling and pigment and metabolic pathways, including melanogenesis, tyrosine metabolism, and retinoic acid biosynthesis. Collectively, these results indicate broad genotype-associated changes in gene expression across neurodevelopmental pathways, setting the stage for detailed investigation of alternative splicing differences between groups.

### Widespread alternative splicing rewiring in HNRNPH2-R114W organoids

A graphical model of the splicing defects in HNRNPH2-R114W organoids is provided in Figure 3A-C. In this model, HNRNPH2 is positioned relative to a spliceosomal assembly (spliceosomal B complex; PDB 8QO9) to demonstrate a plausible spatial relationship between HNRNPH2 RNA engagement within the spliceosomal RNA framework. The HNRNPH2 RNA-binding module is represented by a composite RRM1–RRM3 structure derived from available HNRNPH2 templates listed in the figure legend (PDBs: 1WG5, 1WEZ, 6DG1, 8SGH). At the local interface level, the model highlights the RRM2 component residue R114 at the RNA-recognition surface, positioned within the RRM2–RRM3 region that engages RNA. Based on the short MD refinement and structural superposition with related RNA-recognition antagonist protein templates (SRSF1 and SRSF2; PDBs: 6HPJ, 4C0O, 2LEB), R114 is placed in a geometry compatible with direct hydrogen-bond contacts to nucleotides in the RNA bound to the spliceosome, whereas the R114W substitution is predicted to disrupt this interaction. The precise RNA hairpin or duplex configuration and the global spliceosome docking remain computational predictions that necessitate extended simulations and direct structural validation through methods such as cryo-electron microscopy. However, docking analyses, informed by available HNRNPH2-RRMs 1-3 structures and RNA-recognition protein templates, offer a mechanistic explanation for the manner in which the HNRNPH2-R114W mutation at the RRM2 interface modifies RNA recognition during splicing (Figure 3A-C).

The impact of the HNRNPH2-R114W pathogenic variant on alternative splicing was evaluated using a BAM-based analytical framework, derived from the bulk RNA-Seq datasets. In the junction-centric workflow, splice junctions were quantified with SGSeq, and differential junction usage was assessed using the edgeR quasi-likelihood test. Concurrently, an event-centric exon analysis quantified exon counts with featureCounts (Liao et al., 2014), tested differential exon usage with limma diffSplice (Ritchie et al., 2015), and aggregated gene-level results using the Simes gene-level combining method. Results were summarized as junction-, exon-, and gene-level significance plots, as well as heatmaps of scaled log2 CPM for the most significant differentially used junctions and exons (Fig. 3D-I; Supplemental Fig. 3).

Heatmaps of the top 50 differential junctions and top 37 differentially used exons highlighted a pronounced genotype-dependent clustering (Fig. 3D-E). Plus samples exhibited increased usage of junctions in TUBA1A, TUBB3, CNTN2, and CKB, while minus samples showed higher usage in DCT, PMEL, ALDH1A3, RDH10, CALB1, and XIST. This clear separation indicates a robust splicing signature associated with the HNRNPH2-R114W genotype. Minus-biased exon usage was enriched for calcium and developmental signaling genes, including TRPM3, KNDC1, CDH6, EFS, and the retinoic-acid pathway enzyme RDH10. In contrast, plus-biased exon usage was enriched for neuronal vesicle trafficking, exemplified by VAMP2, cytoskeletal remodeling and actin-associated dynamics, represented by EVL and CRMP1, and RNA or chromatin regulation, represented by HNRNPC, MSI2, and GATAD1.

Junction-level analysis identified over 6,000 significantly altered junctions (FDR < 0.05, |log2FC| ≥ 0.58) out of ∼98,000 tested, representing ∼6–7% of all junctions, shown in a junction-level volcano plot (Fig. 3F), which captures broad, balanced splicing changes, with thousands of junctions regulated in both directions. In contrast, exon-level analysis identified only 17 significant exons (Fig. 3H; Supplemental Fig. 3D-F), indicating that splicing changes predominantly occur at the junction level and are more conservative at the exon level. Gene-level exon usage analysis using a Simes-type framework identified nine genes with significant exon-level changes, including CDH6, EFS, EVL, HNRNPC, UTRN, CRMP1, RUFY3, SPAG9, and PTPRS, which encode proteins involved in neuronal projection, axon development, extracellular matrix organization, and cytoskeletal remodeling (Fig. 3H).

To distinguish between genes with differential expression only (DE only), differential exon usage only (DEU only), or both, we categorized results as follows: 1,966 genes were DE only, seven genes were DEU only, and two genes exhibited both differential expression and exon usage alterations (Supplemental Fig. 3A-C). The seven DEU-only genes, RUFY3, EFS, SPAG9, CRMP1, HNRNPC, UTRN, and EVL, showed significant splicing changes without detectable expression shifts, suggesting regulation primarily at the post-transcriptional level (Supplemental Fig. 2C). CDH6 and PTPRS represent genes with combined transcriptional and isoform-level alterations (Supplemental Fig. 3B).

Concordance between SGSeq and edgeR was evaluated by comparing per-gene significance in overlapping significant genes (Fig. 3I; Supplemental Fig. 3D-F). Genes such as CNTN2, HES5, CYP1B1, COL9A2, IFT172, CLASP1, and TMEFF2 demonstrated consistent significance across both analytical approaches. This agreement between methods reinforces the robustness of genotype-associated splicing differences and suggests that the primary signal is not attributable to a single representation of junction usage, as it is consistently observed in both the SGSeq feature-based and the junction-centric edgeR analyses.

Genome-wide SGSeq feature-level differential usage analysis complemented junction-based findings by identifying significant usage shifts at the feature level (Supplemental Fig. 3D). Within the SGSeq framework, a feature is defined as a local splice-graph element based on observed exon–junction structure, which enables detection of complex usage changes not always captured by a single annotated junction. This approach identified significant features in genes such as VIPR1, BSDC1, ITGAV, SNX17, SELENON, GAD1, and KCNN3. These genes are associated with core cellular programs, as indicated by enrichment terms including vesicle membrane, actin-based cell projection, and inorganic cation transmembrane transport. This pattern is consistent with coordinated changes in membrane trafficking, neurite-associated structural organization, and ion-handling pathways that exhibit genotype-associated differential usage. Overall, these analyses indicate widespread junction-centric splicing remodeling in HNRNPH2-R114W rare variant organoids, alongside a more limited set of exon-level usage changes, thereby providing a multi-layered perspective on splicing heterogeneity.

### Integrated RNA-layer metrics identify priority candidates and functional convergence

To organize results across transcriptional and RNA-processing layers, we implemented an integrated gene-centric prioritization framework combining differential expression (DE), event-level (junction-level), and isoform-level evidence (Fig. 4; Supplemental Fig. 4). Event-level evidence captures differential usage of individual splice junctions, whereas isoform-level evidence reflects changes in the relative abundance of full-length transcript isoforms. For each gene, these RNA-processing signals were combined with transcriptional significance to derive a composite prioritization score summarizing aggregate RNA-layer activity.

**Figure 4.**
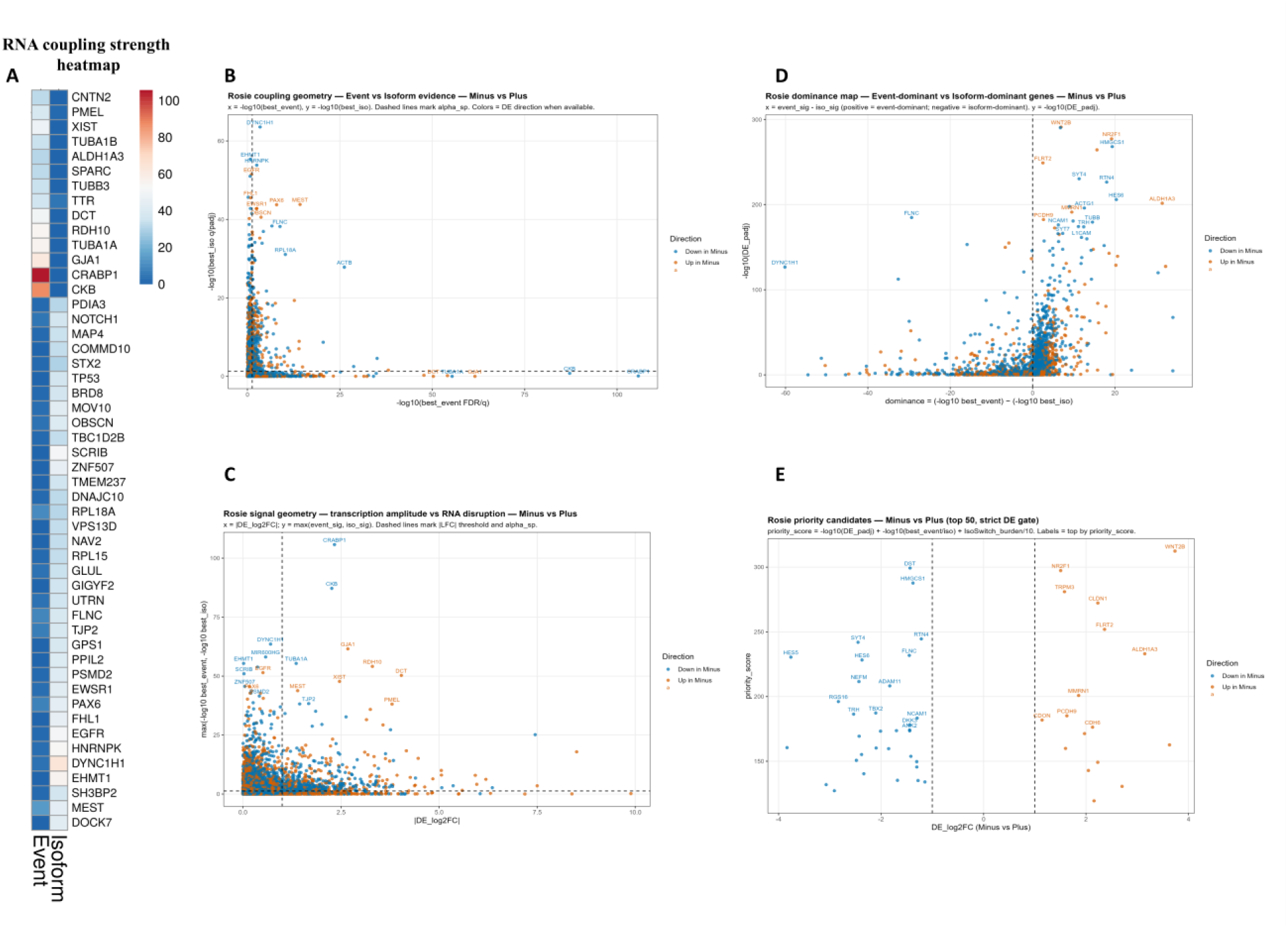
Integrated RNA-layer prioritization connects differential expression with splicing and isoform remodeling, revealing convergent functional programs. (A) The coupling-strength heatmap summarizes integrated multi-layer RNA remodeling signals for prioritized genes, combining gene-level differential expression with event- and isoform-level evidence. (B) Coupling-geometry scatterplot positions genes by event-level versus isoform-level remodeling, distinguishing those dominated by junction or exon events from those dominated by isoform switching. (C) Signal-geometry scatterplot contrasts transcriptional amplitude with RNA disruption, separating expression-dominant from splicing or isoform-dominant effects. Prioritization employs harmonized Ensembl gene identifiers, version-stripped with duplicates collapsed, and applies a uniform integration threshold, where DE is defined as padj < 0.05, baseMean ≥ 10, total read count ≥ 10, and |log2FC| ≥ 1.0, with splicing or isoform significance assessed at FDR < 0.05. Event-level inputs include exon differential usage (diffSplice) and junction differential usage by SGSeq-derived junction usage testing, while isoform-level inputs are aggregated from isoform-focused frameworks implemented in the pipeline. (D–F) Enrichment analyses for prioritized genes are presented across GO Biological Process, GO Molecular Function, and GO Cellular Component as ridge or peak-density panels.

An RNA coupling strength heatmap summarizes, for top-ranked genes, the relative magnitudes of event-level and isoform-level RNA signals (Fig. 4A). The heatmap reveals heterogeneous layer contributions across the prioritized set, highlighting genes with event-dominant, isoform-dominant, and mixed coupling profiles, including CNTN2, PMEL, XIST, TUBA1A, CRABP1, and CKB.

The relationship between event-level and isoform-level evidence was examined using an event-versus-isoform coupling geometry (Fig. 4B), in which genes are plotted by their strongest event-level significance on the x-axis and strongest isoform-level significance on the y-axis. This representation stratifies prioritized genes into regimes with support at both resolutions versus genes driven predominantly by a single layer, with points shifting toward one axis when the corresponding signal dominates. Representative genes labeled in the plot illustrate these coupling regimes, including DYNC1H1, EHMT1, HNRNPK, ACTB, CRABP1, and CKB.

To relate RNA-processing disruption to transcriptional amplitude, a signal geometry analysis maps absolute DE effect size against an RNA-disruption metric defined as the maximum of event- or isoform-level significance for each gene (Fig. 4C). This plot separates genes with large expression changes but modest RNA-processing signals from genes with relatively modest expression changes accompanied by strong RNA-level perturbation, with representative loci illustrating these regimes, including CRABP1, CKB, TUBA1A, GJA1, RDH10, and PMEL.

A dominance map contrasts the relative strength of event-level versus isoform-level evidence by plotting their numerical difference as a dominance score against overall DE significance (Fig. 4C). Positive dominance values indicate event-dominant genes, negative values indicate isoform-dominant genes, and intermediate values reflect a continuum between these extremes. Representative event-dominant genes include HES6, RTN4, and SYT4, whereas FLNC and DYNC1H1 illustrate isoform-dominant profiles, with NCAM1, L1CAM, and TRH occupying intermediate positions.

Candidates were visualized by plotting transcriptional effect sizes against the composite prioritization score (Fig. 4D), highlighting the top-ranked genes under a strict DE gate, defined by a significant DE-adjusted p-value and an absolute log2 fold-change threshold. Representative top-ranked genes highlighted under this gate include TRPM3, WNT2B, NR2F1, CLDN1, FLRT2, ALDH1A3, HMGCS1, DST, and RTN4.

Functional enrichment analysis of prioritized gene sets was performed separately for GO Biological Process, Molecular Function, and Cellular Component (Supplemental Fig. 4B-D). In BP, the strongest signals include neuronal differentiation and projection programs, including axon guidance and neuron projection morphogenesis, alongside translation and peptide biosynthetic processes, and structural programs including extracellular matrix and cilium organization. In MF, enriched terms emphasize cytoskeletal, adhesion-associated binding activities, and RNA binding. In CC, the enriched categories include pigment granule and melanosome compartments, as well as the nucleus, focal adhesion, and collagen-containing extracellular matrix.

### Cell-type composition shifts in HNRNPH2-R114W organoids

To understand the overall cell-type composition shifts in HNRNPH2-R114W organoids, bulk RNA-Seq samples were analyzed using MuSiC deconvolution approach integrated with an HNOCA human neural organoid atlas generating per-sample estimates of cell-type proportions (Wang et al., 2019; He et al., 2024). Both Minus and Plus organoids comprised mixtures of neuroepithelium, dorsal and ventral telencephalic neuronal populations, glioblasts, oligodendrocyte progenitor cells (OPCs), astrocytes, and non-telencephalic neuron compositional markers (Fig. 5A). Genotype-associated compositional differences revealed that neuroepithelium and glial-biased populations (glioblast, OPC) displayed negative delta fractions (CRISPR **−** variant), indicating enrichment in Minus relative to Plus, whereas selected neuronal populations exhibited positive deltas consistent with a modest neuronal reduction in Minus organoids. Across samples, neuronal fractions in Minus organoids were approximately 5% lower on average than in Plus organoids (Fig. 5B).

**Figure 5.**
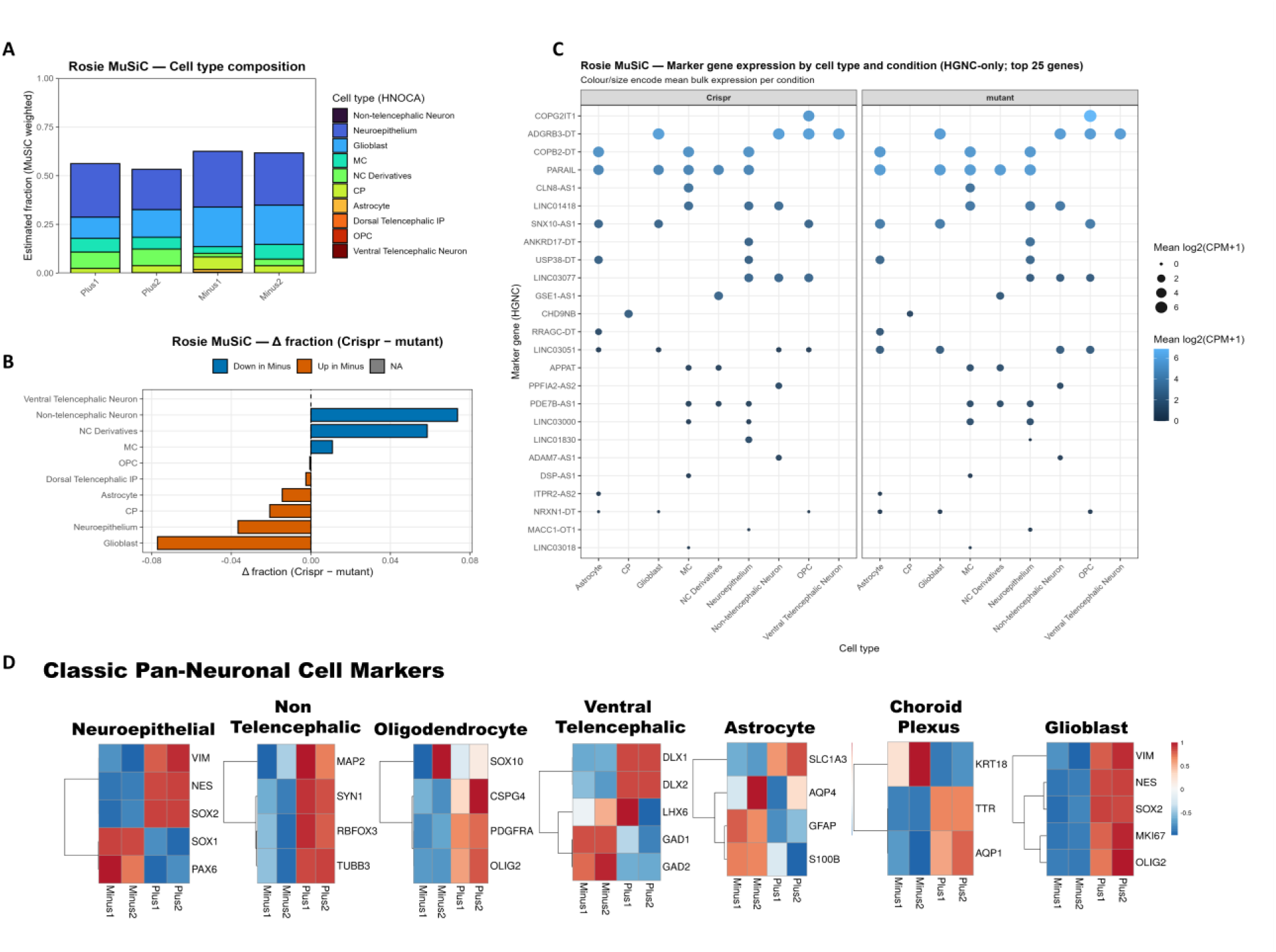
Cell-type composition differences are modest and reproducible by MuSiC deconvolution and marker-program validation. (A) Stacked cell-type fraction plot showing MuSiC-inferred composition per sample for Minus and Plus organoids, estimated by weighted nonnegative least squares regression against a neurodevelopmental single-cell reference (HNOCA) using a shared gene domain (bulk ∩ reference; optionally intersected with MuSiC “used genes”, and retaining cell-type groups supported by a minimum number of five cells per donor. (B) Delta-fraction summary of CRISPR-corrected (plus) versus HNRNPH2 R114W (minus) showing genotype-associated composition shifts across inferred cell types. MuSiC is run on raw (untransformed) gene-level counts, with Ensembl version stripping and duplicate collapsing. (C) Composition concordance assessment using the correlation structure of inferred fractions across samples. (D) Marker validation panel showing conventional marker gene patterns spanning major neuronal cell types; marker genes are computed from log2(CPM+1), centered within each sample, then averaged across markers with z-scaling for visualization.

High within-genotype correlations of MuSiC-inferred fractions indicated stable and reproducible compositional estimates across biological replicates, while lower between-genotype correlations reflected genotype-associated differences rather than random variation (Fig. 5C; Supplemental Fig. 5). These correlation patterns support the robustness of the inferred compositional shifts.

To independently assess whether inferred compositional differences were reflected at the expression level, we examined curated marker gene heatmaps stratified by major lineage classes (Fig. 5D). Neuroepithelial and progenitor-associated markers, including PAX6 and SOX2, astrocyte markers GFAP, S100B, and AQP4, displayed genotype-linked expression patterns consistent with the MuSiC-inferred enrichment of progenitor or glial populations in Minus organoids. Conversely, glioblast markers including MKI67, OLIG2, and the OPC markers CSPG4 and PDGFRA, showed reduced expression in Minus samples, in agreement with the modest reduction in neuronal fractions inferred by deconvolution.

Together, the convergence of MuSiC-based cell-type fraction estimates, lineage-specific marker heatmaps, and expression changes in canonical cell-type markers indicates that HNRNPH2-R114W dysfunction is associated with modest but reproducible compositional biases, favoring neuroepithelial and glioblast lineages over specified neuronal populations. These compositional shifts were reflected by genotype-linked expression changes in canonical lineage markers. Accordingly, although cell-type composition should be considered when interpreting bulk transcriptomic and splicing signatures, the limited magnitude of the compositional differences indicating approximately 5% in neuronal specification reduction suggests that the major gene expression and splicing alterations observed in HNRNPH2-R114W organoids are primarily driven by intrinsic genotype-dependent regulatory effects rather than by changes in cellular composition alone.

## DISCUSSION

### Organoid differentiation as a robust model to investigate the molecular and cellular consequences of the HNRNPH2-R114W mutation

Early corticogenesis constitutes a critical window for dissecting neural developmental processes and informing therapeutic strategies for HNRNPH2 mutations. In this study, isogenic cerebral organoids derived from a patient-specific iPSC line harboring the HNRNPH2-R114W variant (Minus) and its CRISPR-corrected counterpart (Plus) serve as an experimentally rigorous platform for probing RNA-mediated disease mechanisms in a controlled genetic context. Both iPSC lines demonstrate robust expression of canonical pluripotency markers at the stem cell stage and exhibit comparable cellular states prior to differentiation (Fig. 1B). Following the established differentiation protocol, PAX6-positive neural progenitors and TUBB3-positive early neurons consistently emerge in both genotypes (Fig. 1C-D; Supplemental Fig. 1G-H). This indicates that the HNRNPH2-R114W variant does not impair early corticogenesis in vitro. Collectively, these results reinforce the utility of organoid models for recapitulating early neural development in isogenic backgrounds relevant to neurodevelopmental disorders.

HNRNPH2 mutations are causative for an X-linked neurodevelopmental syndrome marked by developmental delay, intellectual disability, Rett- or autism-like features, epilepsy, and variable structural brain abnormalities. These syndromes frequently result from missense mutations within the PY-NLS and qRRM domains (Bain et al., 2016; Pilch et al., 2018; Harmuth et al., 2021; Gillentine et al., 2021). Comprehensive analyses of the heterogeneous nuclear ribonucleoprotein (HNRNP) family reveal that HNRNPH2 variants induce overlapping neurodevelopmental phenotypes, notably including autism spectrum disorder features, by perturbing convergent molecular pathways (Gillentine et al., 2021; Brownmiller & Caplen, 2023). The HNRNP H/H2 proteins specifically bind G-rich RNA sequence motifs, thereby orchestrating splicing regulation, 3′-end processing, and ribonucleoprotein complex assembly (Honoré et al., 1995; Caputi & Zahler, 2001; Huelga et al., 2012; Conboy, 2017; Brownmiller & Caplen, 2023). Studies utilizing patient and neuronal models implicate toxic gain-of-function effects and protein mislocalization as drivers of neural splicing disruptions (Larizza et al., 2022; Brownmiller & Caplen, 2023). The current organoid findings extend these mechanistic models to human cortical tissue and substantiate the interpretation of layered RNA signatures in the setting of RNA-binding protein dysfunction.

### HNRNPH2-R114W genotype defines a global transcriptional state shift

Transcriptomic analysis using DESeq2 reveals that the HNRNPH2-R114W genotype primarily determines transcriptional state, with Minus and Plus samples displaying clear separation and strong intra-genotype concordance. Principal component and Multidimensional analysis further suggest that this separation is cell-intrinsic rather than due to batch or differentiation noise (Fig. 2C; Supplemental Fig. 2) (Love et al., 2014; Qian et al., 2016; Velasco et al., 2019). Broad transcriptional changes identified by differential expression analysis indicate extensive remodeling of global gene expression programs (Fig. 2D–E). More specifically, Plus organoids are enriched for neuronal and synaptic transcripts, while Minus organoids show elevated expression of genes involved in pigment and retinoid metabolism (Fig. 2F-H, Supplemental Fig. 2). Thus, these signatures extend prior studies of HNRNPH2 and HNRNP disorders, reinforcing that HNRNPH2-R114W drives marked, organized transcriptomic reprogramming in cortical organoids (Gillentine et al., 2021; Brownmiller & Caplen, 2023; Larizza et al., 2022; Raj & Blencowe, 2015; Weyn-Vanhentenryck et al., 2018; Irimia et al., 2014).

### Junction-centric splicing rewiring emerges as the dominant molecular phenotype

Splicing analyses link the HNRNPH2-R114W mutation to widespread dysregulation at the junction level. These changes are only partially detected by exon-level tests and cannot be explained solely by gene expression (Fig. 3D-G). Using junction-centric edgeR testing, we identified significant differential usage in about 6–7% of splice junctions, a high proportion for a single RNA-binding protein (Fig. 3F) (Goldstein et al., 2016; Robinson et al., 2010; Brownmiller & Caplen, 2023). Notably, small changes in junction usage can propagate into widespread shifts in isoform abundance across the transcriptome. In contrast, exon-level analysis identified only 17 significant exons in nine genes, highlighting that HNRNPH2 spliceopathy predominantly affects junction usage and isoform balance rather than broad exon skipping (Fig. 3G-H). This observation is consistent with intron- and junction-centric models (Pervouchine et al., 2013; Goldstein et al., 2016).

At both junction- and exon-usage levels, unsupervised heatmaps show pronounced splicing feature separation, especially for junctions (Fig. 3D-E). Genes affected participate in neural development, signaling, and metabolism, supporting pathway-level findings and consistent with HNRNP H-family splicing research (Caputi & Zahler, 2001; Huelga et al., 2012; Ule & Blencowe, 2019; Raj & Blencowe, 2015). The decoupling of junction- and exon-level changes shows substantial RNA-processing pathology can occur without gene-level abundance changes, a pattern seen in other HNRNPH2 and RBP pathology models (Brownmiller & Caplen, 2023; Gillentine et al., 2021; Larizza et al., 2022).

Integrated DE–DEU analysis clarifies intersections between splicing and expression, showing that most genes are DE-only, whereas only seven genes are DEU-only (Supplemental Fig. 2). Specifically, DEU-only genes such as RUFY3, EFS, SPAG9, CRMP1, HNRNPC, UTRN, and EVL demonstrate splicing changes without gene-level expression shifts, which aligns with post-transcriptional regulation. These genes are linked to neuronal wiring and cytoskeletal dynamics, reinforcing themes seen in HNRNP-driven splicing networks and ASD-linked splicing regulators (Irimia et al., 2014; Raj & Blencowe, 2015; Brownmiller & Caplen, 2023). While most transcriptional changes lack detectable exon-level splicing, a smaller subset of genes displays specific or combined disruption, clarifying the relationship between gene expression and splicing alterations.

SGSeq feature-level analyses and cross-method comparisons confirm that major splicing signals are robust across strategies, showing the broad scope of HNRNPH2-dependent RNA rewiring (Fig. 3I; Supplemental Fig. 3). Volcano plots and per-gene overlaps highlight significant events, including shared hits between SGSeq and edgeR (Goldstein et al., 2016; Robinson et al., 2010). This convergence shows that the main targets reflect genuine regulatory changes, consistent with HNRNP proteins’ cooperative regulation of splicing (Huelga et al., 2012; Ule & Blencowe, 2019). Method-specific signals indicate that different analytical approaches capture distinct aspects of RNA architecture, supporting pervasive splicing rewiring during cortical tissue development.

### Multi-layer RNA analysis reveals convergent regulatory hubs

The integrated RNA-layer framework, though showing limited convergence, provides structured and informative insights for disease mechanisms (Fig. 4A-C). Through integrating expression, splicing, and isoform-switch data, priority candidates that intersect transcriptional and post-transcriptional dysregulation are identified (Supplemental Fig. 4) (Love et al., 2014; Robinson et al., 2010; Goldstein et al., 2016; Li et al., 2016; Ule & Blencowe, 2019). For instance, genes such as CRABP1, CKB, TUBA1A, CNTN2, NCAM1, and TRPM3 display multi-layer signatures, whereas others are dominated by isoform-or junction-level changes (Fig. 4B-C; Supplemental Fig. 4). Together, these findings support that RNA-binding proteins affect both broad isoform processing landscapes and more localized RNA splicing events (Caputi & Zahler, 2001; Conboy, 2017).

Top genes such as CNTN2 and NCAM1, crucial for cortical wiring, exhibit multi-layer perturbation and are linked to axon guidance and cell adhesion in cortical development and ASD (Fig. 4A-C) (Irimia et al., 2014; Nowakowski et al., 2017; Pollen et al., 2019), while TRPM3, HES5/HES6, RTN4, and HMGCS1 are involved in neuronal excitability, neurogenesis, and metabolic support. Enrichment analyses highlight pathways in axon guidance, morphogenesis, and synaptic function (Fig. 4D-F). These findings suggest a small set of multi-layer genes function as hubs for HNRNPH2-R114W dependent dysregulation, providing targets for intervention such as antisense oligonucleotides.

### Cell-type composition is a modifier, not a driver, of developmental trajectories

MuSiC deconvolution and pan neuronal-marker analyses reveal that the HNRNPH2 mutation is associated with subtle, reproducible shifts in cell-type proportions rather than lineage turnover (Fig. 5A-D). Minus organoids show higher neuroepithelium and glial populations and reduced neuronal fractions (Fig. 5A-C) (Wang et al., 2019; He et al., 2022). MuSiC modeling and correlation heatmaps confirm these differences are genotype-linked, not artifacts (Fig. 5C). Classic curated marker heatmaps also support the biological plausibility of these shifts (Fig. 5D).

These composition changes suggest HNRNPH2 mutation biases but do not disrupt developmental trajectories. Single-cell atlases show regulated transitions from glia/progenitors to neurons, governed by splicing and transcriptional programs (Nowakowski et al., 2017; Pollen et al., 2019; Quadrato et al., 2017; Pașca et al., 2015). The modest enrichment of progenitor/glial states and reduced neuronal fractions in mutants suggest delayed or altered neurogenic timing, not neuronal loss. This fits clinical data showing microcircuit dysfunction and ASD/ID, not microcephaly (Bain et al., 2016; Pilch et al., 2018; Gillentine et al., 2021; Brownmiller & Caplen, 2023). Overall, the shifts are too small to explain the broad molecular changes observed, supporting cell-intrinsic regulatory dysfunction as the main driver.

Taken together, this work demonstrates an HNRNPH2-R114W variant brain organoid phenotype with preserved early neuronal differentiation, broad but organized expression changes, junction-dominant splicing rewiring, and convergence on key gene hubs, alongside cell composition shifts that act as modifiers, not primary differentiation drivers. These findings support models where HNRNPH2 variants disrupt post-transcriptional regulation, via altered protein localization, splicing, and RNP assembly, producing toxic or dominant-negative effects rather than simple loss of function (Bain et al., 2016; Harmuth et al., 2021; Brownmiller & Caplen, 2023; Gillentine et al., 2021). PY-NLS and qRRM mutations may impair nuclear import and RNA-binding, limiting HNRNPH1 compensation and causing gain-of-function effects in neural splicing networks (Brownmiller & Caplen, 2023; Gillentine et al., 2021). The organoid model provides experimental support for this perspective and offers readouts for allele-selective antisense oligonucleotide strategies or HNRNPH2 reduction to restore paralog compensation (Brownmiller & Caplen, 2023). This integrative approach combining differential expression, multi-layer splicing analysis, and single-cell-based bulk RNA-seq deconvolution should apply to other RBP-driven neurodevelopmental disorders (Ule & Blencowe, 2019; Li et al., 2016; Raj & Blencowe, 2015).

## METHODS

### Derivation and reprogramming of iPSCs

Peripheral blood mononuclear cells (PBMCs) were collected from an individual carrying a pathogenic HNRNPH2 variant. These cells were reprogrammed into induced pluripotent stem cells (iPSCs) using non-integrating episomal vectors under feeder-free conditions by Applied Stem Cell, Inc. An isogenic control line was generated by correcting the HNRNPH2-R114W locus with CRISPR-Cas9 in the same genetic background. The HNRNPH2 rare variant iPSC line was used as the parental line for all subsequent stem cell and organoid differentiation experiments.

### iPSC maintenance culture

iPSCs were maintained on growth factor–reduced Matrigel–coated plates in mTeSR at 37°C, 5% CO2, and greater than 90% relative humidity, with daily medium changes. Cells were passaged using Accutase at 70–80% confluence, reseeded at empirically determined densities to preserve pluripotent morphology, and transiently treated with ROCK inhibitor (Y–27632) following passaging, particularly when dissociated to single cells.

### Pluripotency and genomic QC

Pluripotency of both the HNRNPH2-R114W variant and CRISPR-corrected iPSC lines was evaluated by assessing colony morphology, growth stability, absence of widespread differentiation, and immunostaining for NANOG, SOX2, and OCT4.

### Passage selection for differentiation

To minimize variation from passaging, organoid differentiation was initiated using iPSC cultures at passage 13. Rare variant-carrying and CRISPR-corrected lines were matched for passage number and culture conditions before embryoid body formation. At this stage, colonies showed dense morphology, a high nuclear–cytoplasmic ratio, and minimal edge differentiation. Passage 13 was chosen based on pilot data showing stable pluripotency marker expression between passages 10 and 15. These optimized preparations served as the basis for subsequent cerebral organoid differentiation.

### Cerebral organoid differentiation

Cerebral organoids were generated using a modified Lancaster protocol consisting of four phases: embryoid body formation, neural induction, Matrigel embedding with neuroepithelial expansion, and maturation in suspension culture. For long-term organoid growth, a CO2-resistant orbital shaker was used instead of a spinning bioreactor.

### Wholemount immunostaining and imaging

Organoids were collected at defined developmental time points corresponding to early corticogenesis and cell network establishment. Multiple organoids were harvested per condition and time point. Organoids were rinsed in PBS, fixed in 4% paraformaldehyde for 30–60 minutes at room temperature with mild rocking, washed in PBS to remove fixative, and stored at 4°C in PBS. Fixed organoids were permeabilized in PBS containing 0.2–0.5% Triton X-100 for 1–2 hours at room temperature with gentle agitation, then blocked in PBS with 0.1–0.2% Triton X-100, 5–10% normal serum, and optionally 1–3% BSA for 1–2 hours at room temperature.

### Primary antibody staining

Whole-mount immunostaining was conducted using primary antibodies targeting cortical progenitor and neuronal markers (e.g., PAX6, SOX2, TUBB3), HNRNPH2, and additional study-specific proteins as detailed in the antibody table. Primary antibodies were diluted in blocking buffer at empirically optimized concentrations (typically 1:200–1:1000), applied in sufficient volume to cover organoids, and incubated overnight at 4°C with mild rocking, followed by multiple washes in PBS containing 0.1–0.2% Triton X-100.

### Secondary antibodies and nuclear dyes

Fluorophore-conjugated secondary antibodies (e.g., Alexa Fluor 488, 568, 647) compatible with the host species of the primary antibodies were diluted in blocking buffer and applied for 1–2 hours at room temperature in the dark. Organoids were washed repeatedly in PBS containing detergent. Nuclear counterstaining was performed with DAPI or Hoechst at standard concentrations for 10–20 minutes, followed by additional PBS washes. When necessary, organoids were incubated in a refractive-index-matching solution such as PBS–glycerol prior to mounting.

### Mounting and microscopy

Immunostained organoids were mounted in glass-bottom dishes or chamber slides with a small volume of PBS or mounting medium to prevent movement during imaging. Imaging was performed on a Nikon Eclipse microscope using 2× and 4× objectives and proper filter sets. Multiple fields or z-positions were acquired per organoid to visualize relevant structures. Exposure and illumination settings were kept constant within each experiment.

### Image processing and selection

Images were processed using standard software (FIJI/ImageJ), restricting adjustments to uniform global changes in brightness and contrast across images from a given experiment. Maximum-intensity projections and tiled montages were generated when needed. No non-linear or feature-selective manipulations were applied. Representative images were selected to reflect typical morphology and marker expression, and raw image data were retained for potential quantitative analysis.

### RNA-seq Library Preparation

Total RNA was isolated from each sample (0.375 mL) by addition of TRIzol (1.125 mL; Thermo Fisher). The resulting mixture (1.4 mL) was transferred to a PhaseMaker tube (Thermo Fisher). RNA extraction was performed according to the manufacturer’s protocol, and cleanup was conducted using a RNeasy MinElute Cleanup Kit (QIAGEN). RNA integrity (RIN) was assessed using an Agilent Bioanalyzer. Ribosomal RNA was depleted with a RiboZero Gold (Human/Mouse/Rat) kit (Illumina). RNA libraries were constructed using the NEBNext Small RNA Library Prep Set for Illumina (Multiplex Compatible; Cat #E7330L), following the protocol described in Podnar et al. RNA was fragmented at elevated temperature in buffer to generate fragments averaging 200 nucleotides. Fragments were directionally ligated to 5′ and 3′ adaptors, followed by reverse transcription and PCR amplification. Libraries were sequenced on an Illumina NovaSeq instrument (100-nt single-end reads).

### STAR Alignment for Gene-Level Expression Analyses

Single-end RNA-seq reads were aligned to the human reference genome (GRCh38 primary assembly) using STAR (v2.7.11b; Galaxy EU “RNA STAR” tool) in one-pass, annotation-guided mode. A temporary index was generated from the GRCh38 FASTA and corresponding GTF, with an SA pre-indexing string of 14 and sjdbOverhang set to read length minus one. Alignments were output as coordinate-sorted BAM files with index, including SAM tags NH, HI (one-based), AS, nM, NM, MD, XS, jM, and jI. Library-stranded coverage tracks were produced in bedGraph format and normalized to reads per million (RPM). Key parameters included alignSJoverhangMin = 8 for unannotated junctions, alignSJDBoverhangMin = 1 for annotated junctions, alignIntronMin = 20, alignIntronMax = 1,000,000, outFilterMismatchNoverReadLmax = 0.04, outFilterMultimapNmax = 20, and outSAMmultNmax = 1 to report only the primary alignment per read. Non-canonical splice junctions and chimeric alignments were excluded. STAR outputs included SJ.out.tab, a splice-junction BED file, and per-gene read counts (ReadsPerGene.out.tab) for quality control. Post-run quality control involved review of Log.final.out (unique mapping rate, mismatch rate) and SJ.out.tab motif distributions prior to downstream splicing analyses.

### STAR Alignment for Junction and Exon Splicing Analyses

For splicing-specific analyses, single-end reads were aligned with STAR v2.7.11b in Galaxy EU (“RNA STAR”) using a GRCh38 FASTA and corresponding GTF, index built with GTF; exon features used for splice-junction annotation; SA pre-indexing string length 14; annotated junction sequence length 100. Two-pass mapping and chimeric alignments were not applied. The workflow generated a coordinate-sorted BAM file and splice-junction outputs from STAR, including SJ.out.tab and a corresponding splice_junctions.bed file. BAM alignment tags included NH, HI (one-based), AS, nM, NM, MD, jM, and jI, with the primary alignment flag set to OneBestScore. Unmapped reads and alignments spanning non-canonical splice junctions were excluded using the wrapper’s filtering options, which removed unannotated non-canonical junctions and all non-canonical junction alignments. The unique-mapper MAPQ setting was 255.

### STAR Outputs Used for Downstream Splicing Analysis

BAM: sample.sorted.bam with BAI index

Junction table: SJ.out.tab

Splice junctions BED: splice_junctions.bed

Gene counts (QC): ReadsPerGene.out.tab

Logs: Log.final.out

### Differential gene expression analysis

#### Input data and preprocessing

Gene-level differential expression was performed using DESeq2 on a featureCounts-derived raw count matrix (counts_matrix_FC_stable.csv) containing integer counts per gene and sample. This analysis used organoids generated from matched, quality-controlled batches. Ensembl gene IDs were stored in the first column, sample columns matched entries in a separate metadata table (sample_metadata.csv) containing at least sample and condition fields, sample names were standardized to match column names, Ensembl versions were stripped of duplicate IDs and summed, and genes with fewer than 10 total reads were removed.

#### Group assignment and design

Samples were assigned to Plus (CRISPR-corrected control) and Minus (HNRNPH2-mutant) groups based on controlled metadata (Plus/Control/Crispr versus Minus/Mutant). Samples with ambiguous or unsupported labels were excluded from further analysis. Filtered counts were used to create a DESeqDataSet with the design formula ∼ group.

#### Model fitting and testing

Size factors were estimated using the median-of-ratios method, followed by dispersion estimation and empirical Bayes shrinkage toward a fitted trend. Wald tests were applied to all genes passing the count filter. P-values were adjusted using the Benjamini–Hochberg procedure. Genes with adjusted p-value < 0.05 and baseMean ≥ 10 were considered significant. Log2 fold changes were further shrunk using the ashr method, and both unshrunk and shrunk result tables were exported.

#### Effect size labeling and VST

Genes with |log2FC| ≥ 0.58 (1.5-fold change) were labeled as “Up in Minus” or “Down in Minus” for stratification. A blind variance-stabilizing transformation (VST) was applied to the fitted DESeq2 object. The resulting matrix was used for Pearson correlation, principal component analysis (PCA), and multidimensional scaling (MDS) using the top 500 variable genes.

#### Outputs and visualizations

All quality control outputs, including the VST matrix, similarity matrix, PCA and MDS coordinates, and explained variance tables, were saved to disk. DESeq2 results were exported as complete, significance-filtered, and plot-ready tables. A serialized DESeq2 object (dds.rds) was saved, and visualizations (PCA, MDS, similarity heatmaps, MA plots, volcano plots, DEG bar plots) were generated in a separate module using standardized ggplot2-based settings and exported as high-resolution PNG files.

### Differential splicing and junction usage

#### Strandedness and BAM validation

Library strandedness was determined from STAR ReadsPerGene outputs by comparing forward and reverse strand read counts and classifying libraries as forward-stranded, reverse-stranded, or unstranded based on fixed thresholds. BAM files were checked for the presence of indices and compatibility of chromosome naming with a single GRCh38 GTF, and indices were generated if missing.

#### Exon-level quantification and DEU

Reads overlapping annotated exons were quantified with featureCounts at exon-level resolution, with gene-level grouping disabled, strand specificity enforced according to inferred orientation, and multi-mapping reads excluded. Raw exon counts were stored as a SummarizedExperiment and exported, then converted to a DGEList for analysis with edgeR–limma. Data were normalized by TMM, filtered to retain exons with at least five counts in at least two samples, modeled with voom, and tested for differential exon usage with diffSplice. Exon-level and Simes-aggregated gene-level statistics were produced, with FDR < 0.05 used as the significance cutoff.

#### Junction discovery and testing

Splice junctions were identified and quantified using SGSeq from aligned BAM files and the GRCh38 GTF, capturing both annotated and novel junctions supported by split reads. Low-abundance junctions were filtered using minimum per-junction and total count thresholds. Feature-level logCPM values for junctions and exons were computed with a prior count. Differential usage was tested with limma-trend and empirical Bayes moderation, applying Benjamini–Hochberg FDR correction.

#### Junction significance and gene metrics

Junctions with adjusted p-value < 0.05 were classified as differentially used, and |log2FC| ≥ 0.58 thresholds were applied solely for direction labeling and plotting. All tested junctions, including non-significant ones, were retained in exported tables. Gene-level junction metrics (minimum FDR, number of significant junctions per gene, maximum absolute log2 fold change) were computed and saved as integration-ready outputs.

#### Feature annotation and mapping

Exon and junction features were annotated by genomic overlap with gene models from a cached GRCh38 TxDb. Junction coordinates were parsed into genomic ranges and intersected with gene ranges to assign Ensembl gene identifiers. Ensembl IDs were version-stripped, and gene symbols were added using a unified mapping derived from the same GTF and supplemented as needed. Multi-gene features were assigned the first valid Ensembl ID for deterministic mapping. Annotated feature-level and gene-level tables were exported in RDS and CSV formats, along with scalar-only versions for downstream integration.

#### Universe tracking and QC

At each filtering step, the number of retained junctions and exons was recorded to define the tested feature universe. Separate summaries were generated for total detected, non-zero, lightly filtered, and final filtered features. These universe tables were saved to disk to support transparent enrichment and integration analyses.

#### Docking and 3D Model generation

The model depicted in figure 3A-C was constructed by initially aligning models 1WEZ, 1WG5, and 6DG1 (representing RRM1-3 of HNRNPH2) to 6HPJ and 2M8D (SRSF1) or 2LEB (SRSF2). RRM2 of HNRNPH2 was aligned with 6HPJ or 2LEB, and RRM3 with 2M8D, forming the principal RNA recognition motif using biopython alignment and rigid body modules. Non-modeled regions were subsequently aligned with Swiss-Model, yielding a final model comprising amino acids 5-368. Energy minimization was performed using Gromacs 2024.3 with a steepest descent algorithm, incorporating the RNA from model 6HPJ. The final model was then aligned to 8QO9 to determine the relative position of HNRNPH2 within the spliceosomal B complex. RRM2 again aligned with the lowest RMS score to SRSF1, with a difference of less than 0.5 for main chain atoms. This configuration positioned the RNA to enter the spliceosomal complex directly and demonstrated that R114 was in direct contact with any bound RNA ligand (Waterhouse, et al., 2018).

### Cell-type deconvolution (MuSiC)

#### Bulk RNA-seq input preparation

Bulk RNA-seq inputs for MuSiC were derived from the same featureCounts gene-level count matrix used for DESeq2, using raw counts without transformation to preserve linearity. Ensembl gene IDs were version-stripped, duplicates collapsed by summing counts, library sizes were computed for all samples, and sample metadata were harmonized to ensure one-to-one correspondence between count columns and annotations, including standardized condition labels for mutant (patient) and corrected groups.

#### Single-cell reference and gene domain

A human neurodevelopmental single-cell reference (HNOCA) was used, retaining only cell-type groups with a minimum number of cells per donor to ensure robust pseudobulk estimates. A shared gene domain was defined as the intersection of genes detected in both bulk and single-cell reference matrices and, when available, further intersected with internal “used gene” set to restrict deconvolution to informative genes MuSiC.

#### MuSiC model fitting

Cell-type proportions were estimated in MuSiC using weighted non-negative least squares regression, modeling each bulk sample as a linear combination of reference cell-type expression profiles. All non-zero cell types across samples were retained, estimated fractions per sample were constrained to sum to one, and sample-wise correlation matrices of cell-type fractions were computed to evaluate internal consistency and replicate concordance.

#### Stability and identifiability checks

Regression stability measures were derived from MuSiC gene-level weight matrices, with a per-sample stability score defined as the sum of absolute gene weights. Stability scores were inspected across samples, and relationships between cell-type fractions and stability were examined to confirm that inferred compositional shifts were not driven by numerical instability or model collapse.

#### Marker genes and program scores

Cell-type marker genes were identified from the single-cell reference by computing log-fold changes for each cell type against all others within the MuSiC gene domain, retaining top markers with sufficient foreground and background representation. Bulk marker program scores were calculated by transforming counts to log2(CPM+1), centering within samples, and averaging marker gene expression per cell type; scores were optionally z-scaled across samples for visualization, while raw values were preserved for integration.

#### Deconvolution outputs and visualization

MuSiC outputs, including cell-type proportion tables, regression stability measures, marker gene lists, and program scores, were exported in standardized, integration-ready formats with Ensembl IDs and HGNC symbols. Visualization scripts generated stacked bar plots of cell-type composition, delta-fraction plots comparing conditions, and heatmaps, dot plots, or scatter plots of marker expression against inferred fractions using fixed dimensions and consistent color schemes.

### Pan-layer integration and synthesis

#### Gene-level master tables

Gene-level summary tables were generated for each analytical domain and merged using version-stripped Ensembl IDs as the primary key. DESeq2 differential expression metrics formed the anchor layer, and event-level splicing (diffSplice exon usage and junction usage) and isoform-level remodeling metrics (SGSeq, DRIMSeq, IsoformSwitchAnalyzeR) were aggregated to the gene level and joined to the expression table, with gene symbols harmonized across layers, and Ensembl IDs retained as fallback identifiers.

#### Quality gating and universe definition

A uniform quality gate was applied across layers: genes were considered differentially expressed if adjusted p-value < 0.05, baseMean ≥ 10, total read count ≥ 10, and |log2FC| ≥ 1.0. Splicing significance was assessed separately for event-level and isoform-level analyses using FDR < 0.05, and the DESeq2 gene set defined the reference universe for all downstream analyses, with QC summaries documenting universe size, layer coverage, and gate concordance.

#### Pan-RNA tier assignment

For each gene, binary indicators captured differential expression, event-level splicing disruption, and isoform-level remodeling, enabling assignment to mechanistic tiers: pan-RNA (expression plus splicing and/or isoform changes), event-only (splicing-driven), isoform-only (isoform program–driven), or no detectable RNA-level disruption. When a pre-computed membership table was unavailable, tier assignments were derived on the fly from the evidence flags.

#### Quantitative prioritization

Continuous significance metrics were retained alongside binary indicators, propagating the most significant event-level and isoform-level p-values per gene and converting them to −log10 scores. These scores were combined with DESeq2 effect sizes and significance, and standardized direction labels (“Up in Minus”, “Down in Minus”) were applied to generate ranked gene lists within and across tiers.

#### Gene sets and enrichment analysis

Tier-specific and direction-specific gene sets were exported as TSV files and plain-text lists and subjected to over-representation analysis using Enrichr with Gene Ontology Biological Process, Molecular Function, and Cellular Component libraries. Enrichment analyses used the DESeq2-defined gene universe as the background and enforced minimum gene-set and universe-size thresholds, with results summarized in tables and visualizations highlighting convergent biological themes across tiers.

#### Reproducibility and final outputs

All integration steps were executed in scripted pipelines with automatic column detection, override checks, and comprehensive logging of run parameters, session information, QC metrics, and file inventories. Final outputs included gene-level master tables for each integration axis, pan-RNA membership summaries, tier-specific gene sets, enrichment tables, and publication-ready figures supporting the mechanistic interpretation of RNA-level changes in HNRNPH2-mutant brain organoids.

#### AI-assisted code development and validation procedures

RStudio analysis scripts were developed with assistance from several large language models, including ChatGPT, Claude, and Perplexity, to facilitate code drafting, debugging, and testing. All scripts and outputs were independently cross-checked, reviewed, and validated prior to implementation.

#### Primary antibodies

POU5F1 (OCT4): DSHB, PCRP-POU5F1-1D2 (RRID: AB_2618968).

NANOG: DSHB, PCRP-NANOGP1-2D8 (RRID: AB_2722264).

SOX2: DSHB, PCRP-SOX2-1B3 (RRID: AB_2722343).

PAX6: DSHB, PAX6 (RRID: AB_528427).

TUJ1: DSHB, 2H3 (RRID: AB_531793).

SHANK1: DSHB, N22/21. N22/21 was deposited to the DSHB by UC Davis/NIH NeuroMab Facility (DSHB Hybridoma Product N22/21).

G3BP1: DSHB, PCRP-G3BP1-2H8 (RRID: AB_2722179).

HNRNPH2: Abcam, Anti-HNRPH2/HNRNPH2 antibody [EPR12170(B)] ab179439.

## DATA AVAILABILITY

Bulk RNA-sequencing data from this study are available in NCBI under BioProject PRJNA1413422 at https://dataview.ncbi.nlm.nih.gov/object/PRJNA1413422?reviewer=vfbmrb076l69qri0m0li2pnboh.

## CODE AVAILABILITY

The analysis code used in this study is available at https://github.com/carlodonato/HNRNPH2-Brain-Organoids.

## Institutional Biosafety Compliance and Authorization for NCBI Data Deposition

All stem cell and organoid research was conducted in accordance with institutional biosafety protocols at The University of Texas at Austin, under Institutional Biosafety Committee (IBC) Protocol ID IBC-2023-00276, approved through December 11, 2026. Authorization was obtained to deposit sequencing datasets at the National Center for Biotechnology Information (NCBI). The BioProject and associated Sequence Read Archive (SRA) metadata are accessible to reviewers in read-only format via the NCBI Dataview reviewer link for PRJNA1413422. This access will remain active and reflect all associated metadata until public release.

## Acknowledgements

The authors thank the Brazilian National Council for Scientific and Technological Development (CNPq) for funding Caiaffa, C.D. during manuscript preparation. The authors also thank the To Cure a Rose Foundation for support.

## Author contributions

**Carlo Donato Caiaffa:** Conceptualization, Data curation, Formal analysis, Funding acquisition, Investigation, Methodology, Project administration, Resources, Validation, Visualization, Writing – original draft, Writing – review & editing. **Neda Ghousifan:** Resources, Visualization. **Stephan Lloyd Watkins:** Formal analysis, Methodology, Validation, Visualization. **Helder Nakaya:** Formal analysis, Visualization. **Rodney Bowling:** Conceptualization, Funding acquisition, Project administration, Resources, Visualization. **Richard Finnell:** Conceptualization, Funding acquisition, Project administration, Supervision, Visualization, Writing – review & editing. **Robert Cabrera:** Conceptualization, Funding acquisition, Project administration, Supervision, Visualization, Writing – review & editing.

