## Supplemental Fig. 1-5 for "Decoding the RNA Splicing Network in HNRNPH2-R114W Brain Organoids"

**SUPPLEMENTAL FILE:**

#### Supplemental Figure 1

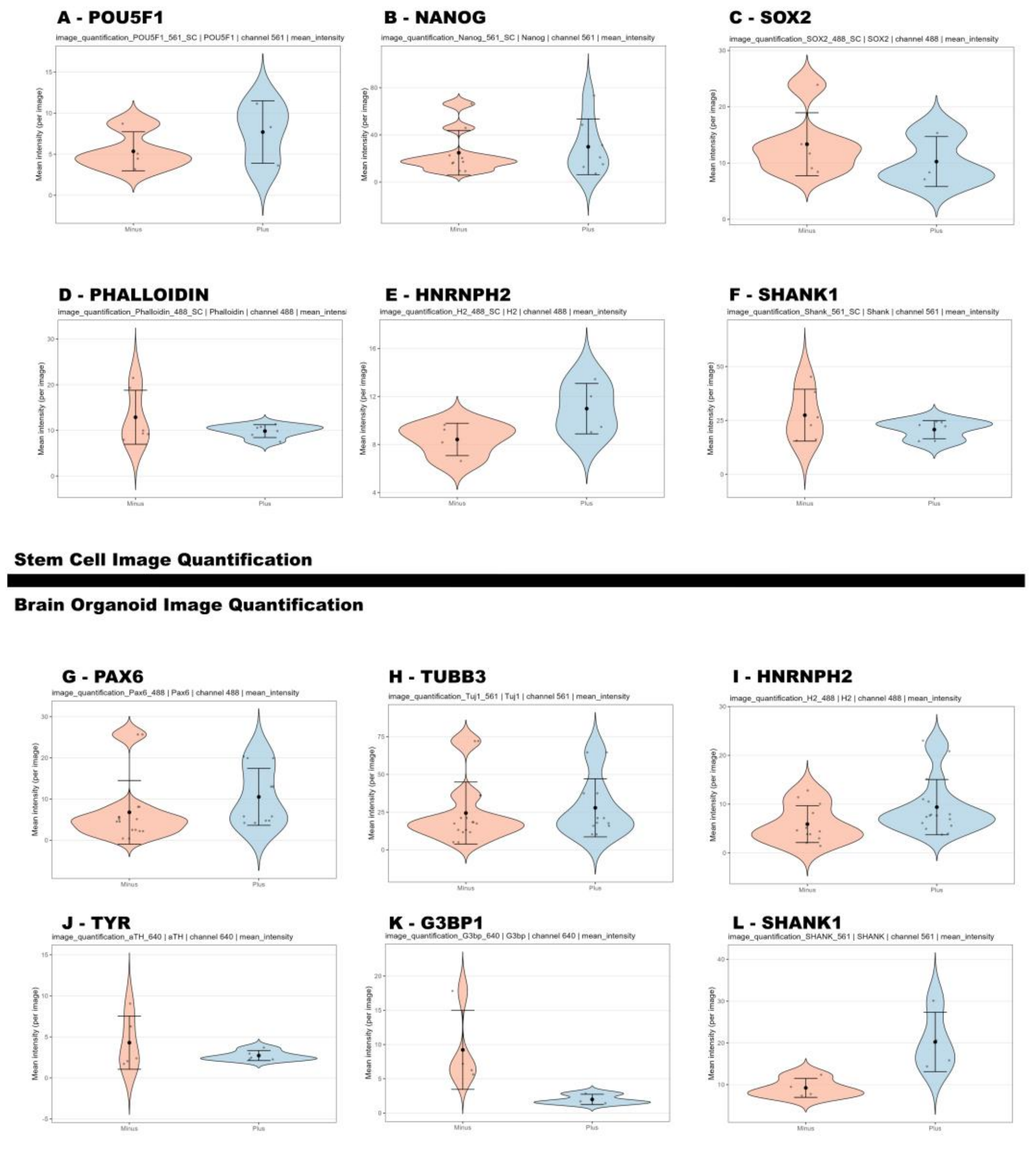


**Supplemental Figure 1. Quantitative immunofluorescence demonstrates preserved pluripotency and early neuroepithelial marker profiles, with distinct lineage-associated marker differences observed between Minus and Plus.** (A–L) Violin plots of per-image mean intensity compare Minus and Plus for iPSC markers (A) POU5F1, (B) NANOG, (C) SOX2, (D) Phalloidin, (E) HNRNPH2, and (F) SHANK1, as well as organoid lineage-associated markers (G) PAX6, (H) TUBB3/Tuj1, (I) HNRNPH2,

(J) aTH/TYR, (K) G3BP1, and (L) SHANK1. Quantification was performed using microscope-exported images filtered by expected channel identifiers in filenames (488, 561, 640), restricting analysis to files with the corresponding channel. Pixel intensities were extracted from each selected image and summarized per image (mean, sum, and pixel count). Violin plots display the distribution of per-image mean intensity by condition; dots represent individual images, the black point indicates the group mean, and the vertical bar represents ±SD.

#### Supplemental Figure 2

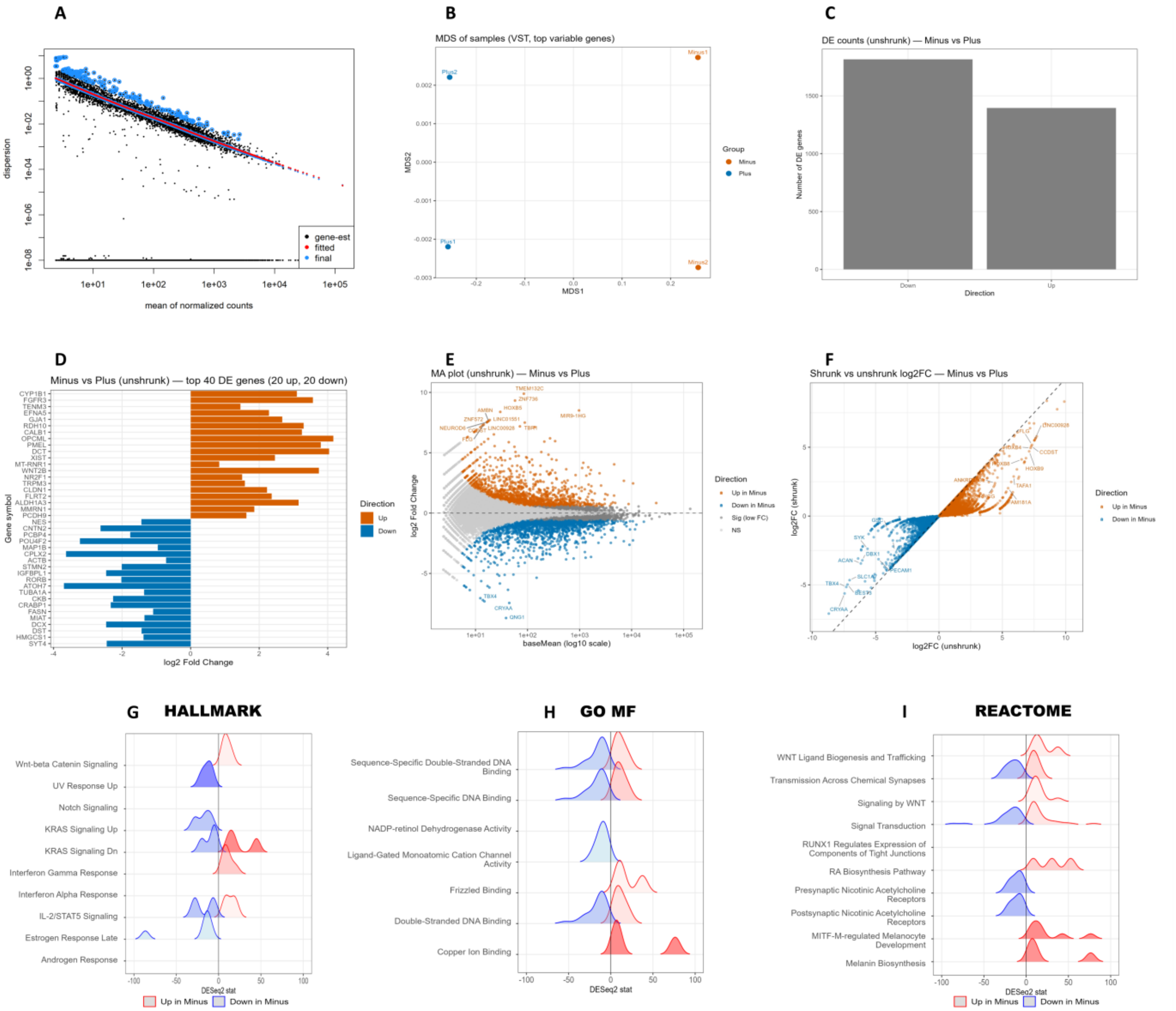


**Supplemental Figure 2. Quality control and effect-size diagnostics reveal clear transcriptomic separation between HNRNPH2 R114W (Minus) and isogenic CRISPR-corrected (Plus), with broad differential expression and direction-resolved enrichment patterns.** (A) DESeq2 dispersion estimates are plotted against the mean of normalized counts (baseMean) with the fitted trend. (B) Multidimensional scaling (MDS) is performed on variance-stabilized, high-variance genes. (C) Bar plot displays the number of significantly differentially expressed genes in each direction using unshrunk log2 fold-change estimates. (D) Ranked bar plot presents unshrunk log2 fold changes for the top 20 upregulated and top 20 downregulated genes in Minus versus Plus. (E) Unshrunk MA plot illustrates log2 fold change as a function of mean expression (baseMean) on a logarithmic scale. (F) Scatter plot compares unshrunk versus shrunk log2 fold changes for all genes. (G–I) Ridge-style enrichment distributions are separated by direction (up in Minus versus down in Minus) for (G) HALLMARK gene sets, (H) GO Molecular Function, and (I) Reactome pathways.

#### Supplemental Figure 3

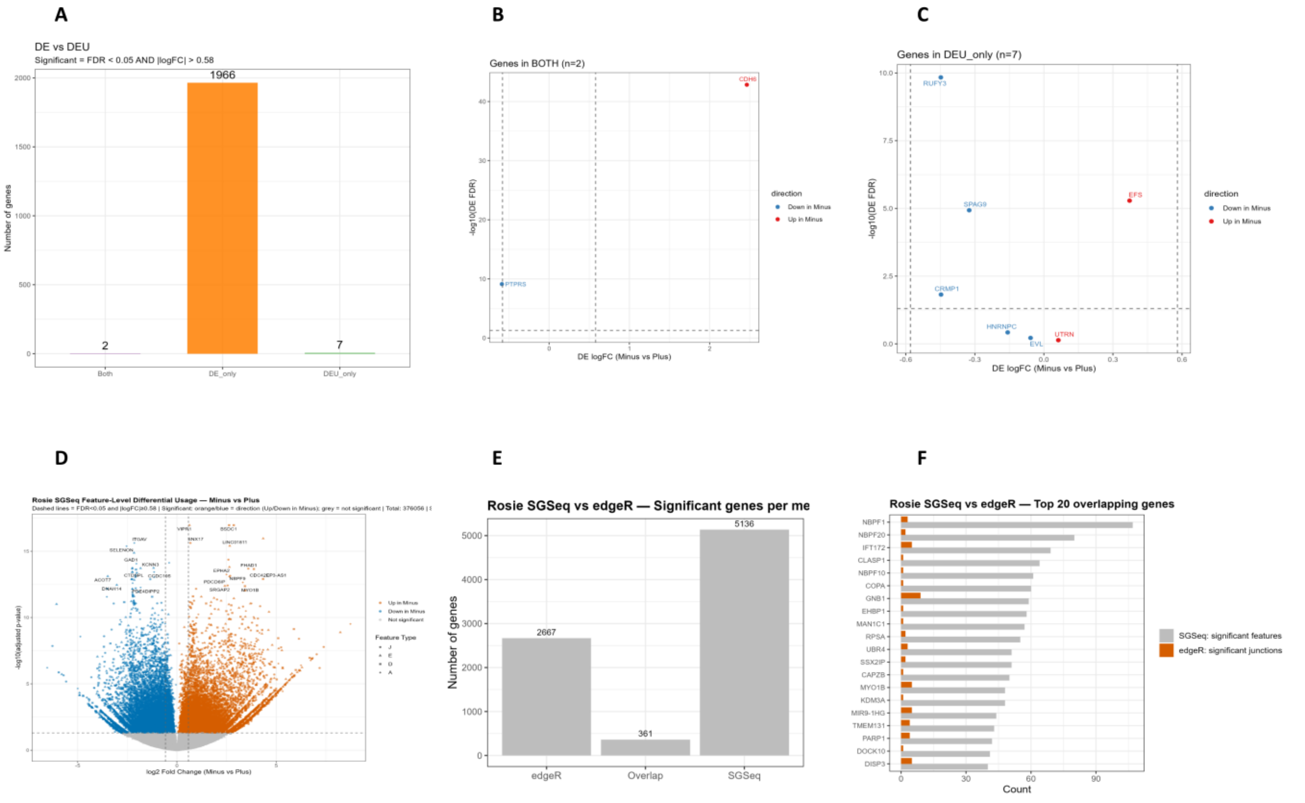


**Supplemental Figure 3. Differential expression and exon-usage categorization, feature-level differential usage, and cross-method concordance collectively summarize splicing-associated signals identified by SGSeq and edgeR.** (A) Bar plot summarizing gene-set categories from the comparison of differential expression (DE) and differential exon usage (DEU) ("DE only," "DEU only," "Both"), with counts for each category under the specified thresholds. (B) Scatter plot of genes classified as "Both" (n=2), displaying DE log2FC (Minus vs Plus) versus DEU signal (−log10 FDR), with point colors indicating direction. (C) Scatter plot of genes classified as "DEU only" (n=7), displaying DE log2FC (Minus vs Plus) versus DEU signal (−log10 FDR), with point colors indicating direction. (D) SGSeq feature-level differential usage volcano plot (Minus vs Plus), displaying log2 fold change against

−log10(FDR) for splice features. (E) Bar chart comparing the number of significant genes detected by each method (edgeR, overlap, SGSeq). (F) Horizontal bar chart of the top 20 overlapping genes, listing the top overlapping targets and method-specific counts (SGSeq significant features versus edgeR significant junctions).

#### Supplemental Figure 4

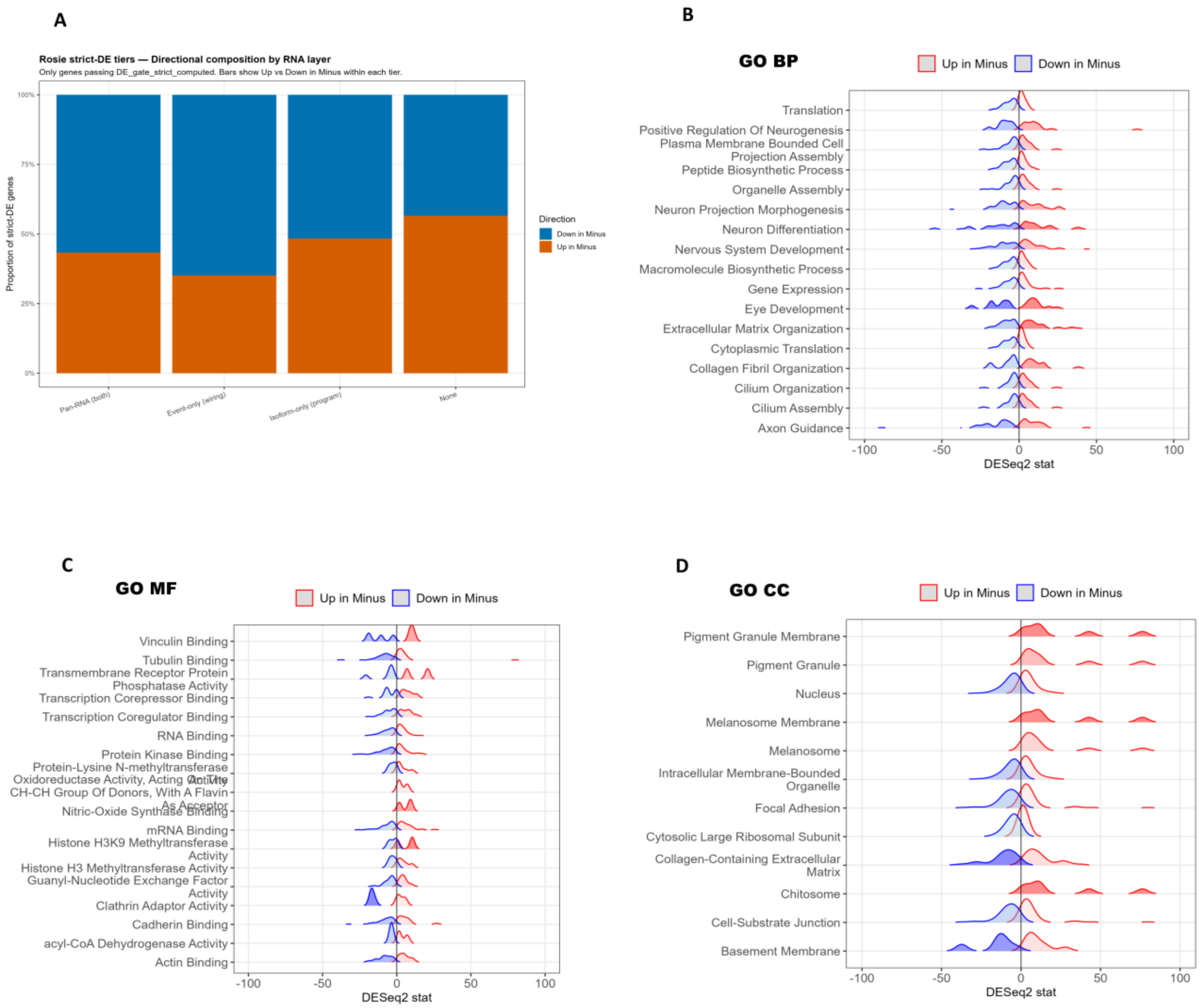


**Supplemental Figure 4. Directional composition by RNA layer and direction-resolved Gene Ontology (GO) enrichment distributions summarize strict differential regulation across functional categories.** (A) Stacked bar plot illustrating the directional composition of strictly differentially regulated genes by RNA layer, with proportions by direction (up in Minus versus down in Minus) across RNA-layer categories. (B) Ridge-style enrichment distributions for GO Biological Process (GO BP), separated by direction (up in Minus versus down in Minus). (C) Ridge-style enrichment distributions for GO

Molecular Function (GO MF), separated by direction (up in Minus versus down in Minus). (D) Ridge-style enrichment distributions for GO Cellular Component (GO CC), separated by direction (up in Minus versus down in Minus).

### Supplemental Figure 5

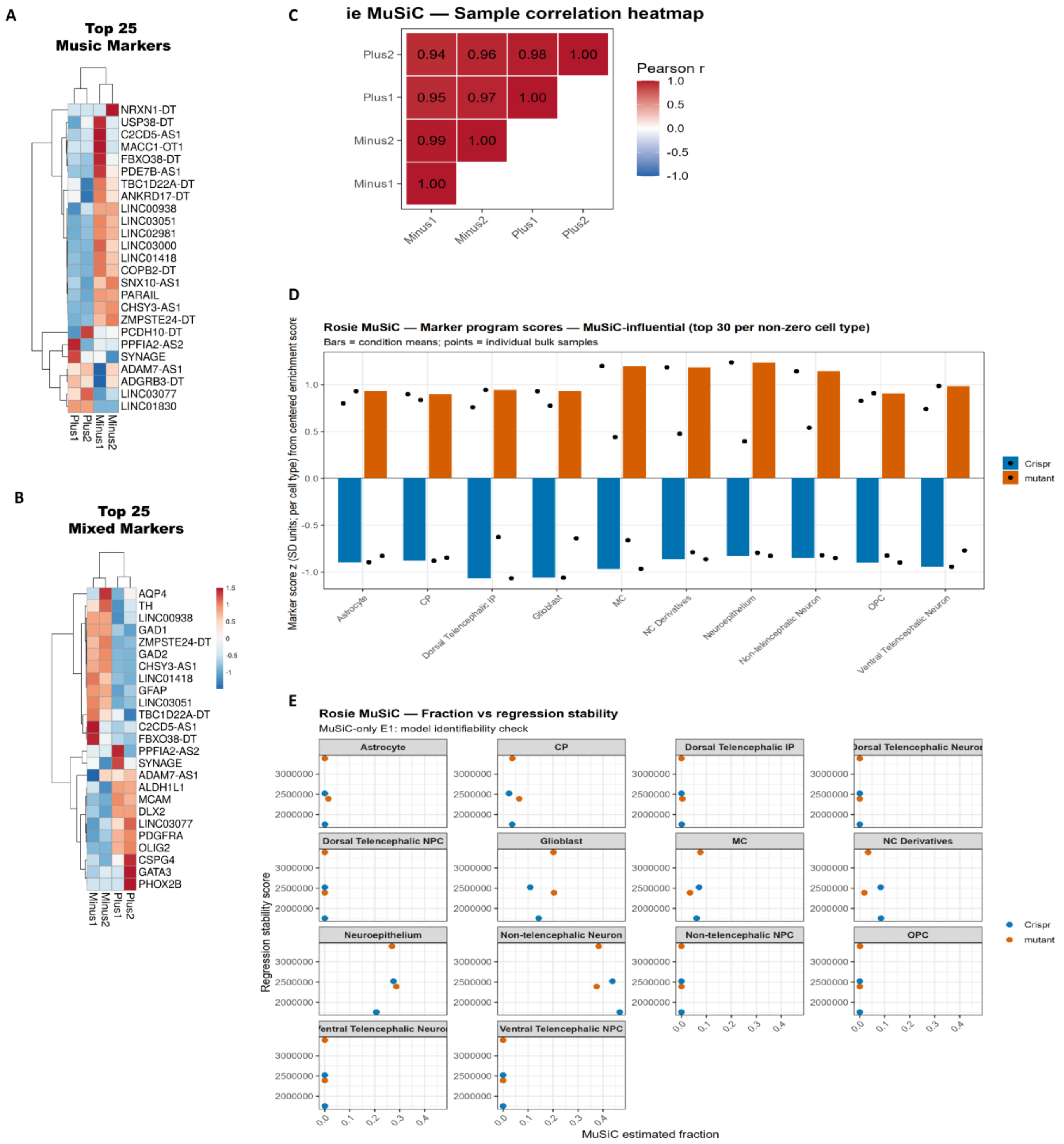

**Supplemental Figure 5. Marker-expression heatmaps, fraction correlation structure, marker program scores, and stability diagnostics collectively summarize MuSiC deconvolution support for cell-type composition patterns.** (A) Heatmap of bulk expression for the "Top 25 MuSiC markers" gene set (rows represent genes with hierarchical clustering and scaled values displayed. (B) Heatmap of bulk expression for the "Top 25 mixed markers" gene set with hierarchical clustering and scaled values displayed. (C) Sample correlation heatmap displaying pairwise Pearson correlations between samples based on MuSiC-derived cell-type proportion identifiers. (D) Marker program score bar plot for MuSiC-influential markers, with bars representing condition means and points representing individual bulk samples, grouped by cell type. (E) Fraction versus regression stability scatter plots, faceted by cell type, displaying regression stability metric against MuSiC estimated fraction per sample, with points colored by condition.
